# Stromal cells modulate innate immune cell phenotype and function in colorectal cancer via the Sialic acid/Siglec axis

**DOI:** 10.1101/2025.03.14.642985

**Authors:** Aoise O’Neill, Norashikin Zakaria, Hannah Egan, Shania M Corry, Courtney Bull, Niamh A Leonard, Clodagh O’Meara, Linda Howard, Anastasija Walsh, Eileen Reidy, Jenny Che, Li Peng, Lizhi Cao, Laurence J Egan, Thomas Ritter, Margaret Sheehan, Aoife Canney, Kevin Culligan, Aisling M Hogan, Sean O Hynes, Philip D Dunne, Michael O’Dwyer, Oliver Treacy, Aideen E Ryan

## Abstract

**Background:** The immunosuppressive tumour microenvironment (TME) reduces immune response effectiveness in stromal-rich tumours, including CMS4 colorectal cancer (CRC). Mesenchymal stromal cells (MSCs), precursors to cancer-associated fibroblasts (CAFs), promote cancer progression by suppressing anti-tumour immune responses. Hypersialylation of glycans on tumours engages Siglec receptors on immune cells, driving immune dysfunction, but its role in stromal-mediated immunosuppression remains unclear.

**Methods:** Sialic acids and Siglec ligands were measured on CRC tissue, primary human CAFs, and tumour-conditioned-mesenchymal stromal cells (MSC^TCS^) and CAF using immunohistochemistry and flow cytometry. The effect of stromal cell sialylation on macrophages and NK cells was assessed in *ex vivo* primary stromal and immune cell co-cultures and expression of Siglec-10 and immune cell phenotype markers and function were measured by flow cytometry. Using an immunocompetent Balb/c CT26 mouse model, we induced tumours with/without conditioned stromal cells, with/without pre-treatment of stromal cells with sialyltransferase inhibitor (3FAX) or sialidase (E610). We assessed the effect of stromal cell sialylation on macrophages and NK cells in the tumour and secondary lymphoid tissues by flow cytometry.

**Results:** Stromal cells, including CAFs, in CRC tumours are highly sialylated compared to epithelial cancer cells and are associated with high expression of *ST6GalNAC6*. Genetic knockdown of *ST6GalNAC6* reduced the expression of stromal cell Siglec-10 ligands in MSCs. CAFs and MSC^TCS^ induced Siglec-10 on macrophages and NK cells and impaired NK cell cytotoxicity. Sialidase treatment reduced Siglec-10 expression, restoring NK cell function. *In vivo*, desialylation of stromal cells increased macrophage activation (CD11b+CD80+) and reduced immunosuppressive marker expression (CD206, PD-L1, Siglec-G) in lymphoid tissues, indicating sustained systemic anti-tumour immunity. Intratumoural NK cells exhibited high Siglec-G expression and impaired cytotoxicity, and granzyme B expression significantly increased with sialidase treatment of stromal cells. In an inflammatory tumour model, inflammatory tumour-conditioned (iTCS) MSCs promoted metastasis and Siglec-G induction on NK cells and macrophages, both reversed by sialyltransferase inhibition, underscoring the effects of stromal modulation of innate immune cell function in inflammatory tumours.

**Conclusion:** Stromal cell sialylation modulates innate immune suppression in CRC via the sialic acid/Siglec axis. Targeting stromal sialylation restores NK cytotoxicity and macrophage activation, offering a novel therapeutic strategy for immunosuppressive stromal-rich tumours.

**What is already known on this topic:**

- The tumour microenvironment of consensus molecular subtype 4 (CMS4) colorectal cancer (CRC) is associated with high stromal burden, poor immune infiltration, poor response to anti-cancer therapies and thus poor patient prognosis. Immune checkpoint inhibitors (ICIs) have limited impact on stromal-rich CRC tumours, therefore highlighting the need to discover and target novel mechanisms of tumour immune evasion.
- Emerging studies have highlighted that stromal cells in CRC and pancreatic ductal adenocarcinoma (PDAC) are highly sialylated, expressing even higher levels of sialic acid on their cell surface than epithelial cancer cells. Targeting stromal cell sialylation has unveiled promising data in restoring the anti-tumour activity of T cells and macrophages. There is a need to explore the effects of targeting stromal cell sialylation on other immune cells of the TME and to evaluate the Siglec/sialic acid axis of stromal and immune cells in resistant CRC tumours.

**What this study adds:**

- We reveal *ST6GalNAC6* as a sialyltransferase enzyme that regulates the production of Siglec-10 ligands in CRC stromal cells. Overexpression of *ST6GalNAC6* and Siglec-10 correlated with poor survival in CRC and mesenchymal CRC tumours.
- We show for the first time an induction of Siglec-10 expression on macrophages and NK cells in stromal-immune co-culture experimental models with hypersialylated MSCs and CAFs *in vitro* and *ex vivo*. Targeting stromal cell sialylation increased NK cell cytotoxicity of CRC cells, indicating a direct functional role for stromal cell sialylation in immunosuppression.
- An immunogenic mouse model of CRC was used to evaluate the potential therapeutic efficacy of targeting stromal cell sialylation in overcoming stromal cell-mediated immunosuppression in CRC. Sialic acid-targeting of stroma slowed tumour growth and reduced inflammation-driven metastasis. This was associated with greater infiltration and activation of macrophages and NK cells with stromal cell sialic acid depletion, highlighting stromal cell sialylation as a mechanism of innate immune cell suppression in stromal-rich CRC.

**How this study might affect Research, Practice or Policy:**

- Our research provides insight into a novel mechanism of stromal cell-mediated immunosuppression of innate immune cells in CRC and may open up new avenues of research for targeting stromal cells in stromal-rich TMEs such as pancreatic, breast and ovarian cancers.
- Our research identifies a stromal cell effect of enhancing Siglec expression on tumour infiltrating innate immune cells as a novel immune checkpoint, which may be useful in identifying potential novel immunotherapeutic combinations in future.

**Graphical abstract:** 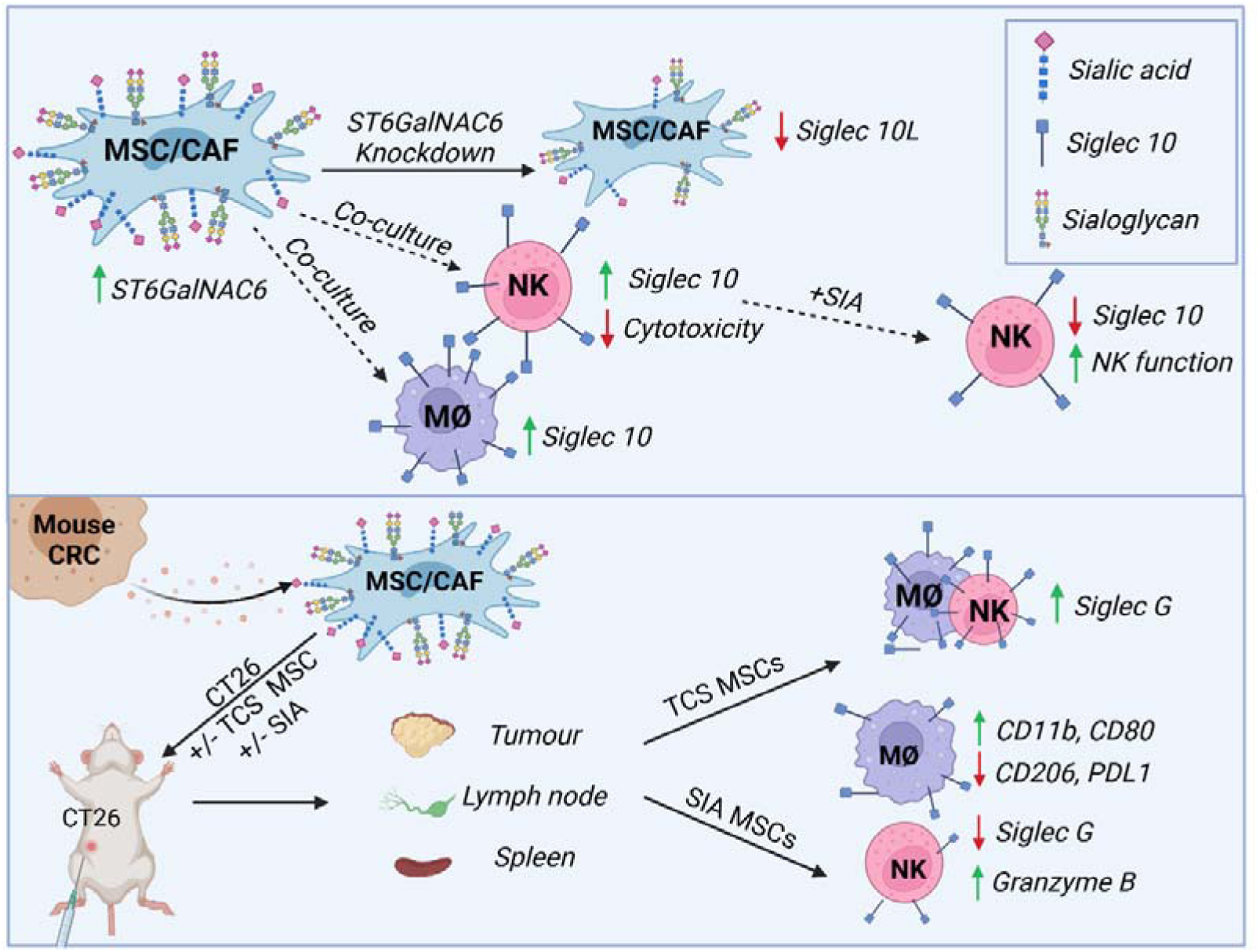

## Introduction

Colorectal cancer (CRC) is the second leading cause of cancer mortality worldwide, primarily due to late-stage diagnosis and resistance to current therapies [1]. Advances in immunotherapies for CRC have been largely unsuccessful, except for microsatellite instable (MSI) tumours, which represent only 15% of all CRC. In microsatellite stable (MSS) tumours, immunosuppression within the tumour microenvironment (TME) is a major therapeutic challenge [2]. CRC can be classified into four consensus molecular subtypes (CMS) based on transcriptional and mutational profiles [3]. CMS1 is associated with high immune cell infiltration, BRAF mutations and good response to therapy, largely due to its enrichment in MSI tumours [4]. Conversely, mesenchymal subtype CMS4 CRC, which has high stromal burden and poor prognosis, is enriched with MSS tumours [5]. Enrichment for MSS tumours and the presence of stromal cells and inflammatory signatures are associated with the worst disease-free survival rates in CMS4 CRC [6]. Therefore, novel therapeutic approaches are urgently needed.

The CMS4 TME is highly complex and heterogeneous, comprising of cancer cells, stromal cells, immune cells, extracellular matrix (ECM), and secreted factors. Stromal cells of mesenchymal origin, which include mesenchymal stromal cells (MSC) and cancer-associated fibroblasts (CAF), predominate the CRC TME and expansion of these cells promotes tumour growth, angiogenesis, metastasis and drug resistance [7].There is growing evidence that their ability to potently induce immunosuppression is a predominant and targetable mechanism of tumour promotion [8–14] and the immunomodulatory properties of stromal cells have been well established [15, 16]. Although stromal cells are a desirable target for immunotherapy, their inherent heterogeneity makes targeting difficult, as MSCs and CAFs have limited specific markers and share many similarities with normal fibroblasts [17]. Therefore, identifying predominant mechanisms of action in tumour promotion, such as those that induce immunosuppression, are of significant interest and may hold promise to reinvigorate anti-tumour immunity and improve outcomes in CRC.

The cancer-immunity cycle now includes the impact of stromal cells in shaping the immunosuppressive landscape in the TME, including the CRC TME [18]. CD8+ cytotoxic T cells have been shown to reside in the stroma of CRC preventing them from infiltrating and killing tumour cells [19]. Macrophages in CRC have been shown to be polarized to an anti-inflammatory, pro-tumorigenic phenotype [20]. Natural killer (NK) cells have reduced cytotoxicity of cancer cells in the presence of stromal cells [9]. The mechanisms of stromal-immune modulation have been of significant interest in recent years [21–23]. Recent evidence from our group and others suggests that post-translational sialylation may be important in regulating stromal cell immunoregulatory functions [19, 24].

Sialylation, a post translational process involving sialic acid addition to glycoproteins and glycolipids, plays a vital role in cellular communication and immune modulation [25, 26]. The biosynthesis and degradation of sialylated ligands is regulated by the expression of sialyltransferases (ST) and sialidases, respectively [27]. Hypersialylation has been linked to immune evasion in many cancers including breast, lung, pancreatic and multiple myeloma [26]. Binding of sialylated ligands to sialic-acid binding immunoglobulin-like lectins (Siglecs) can impair immune function [26]. There are fourteen human Siglecs, Siglec 1-15, which are expressed differentially on innate and adaptive immune cells, and 3 mouse orthologues Siglec-E,-G and-F [28, 29]. Once bound by sialic acid, Siglecs have a downstream ITIM/ITAM signalling motif which inhibits or activates immune responses via SHP1/2 or through SYK, respectively [29]. Siglecs-7,-9 and-10 are the most well characterized Siglecs and are expressed on T cells, NK cells and macrophages [30] [31, 32]. Tumour and stromal cells in the TME can exploit this pathway to evade immune surveillance by expressing Siglec ligands [33, 34]. Some Siglec ligands (Siglec-L) have also been discovered in recent years in the TME and include PSGL-1, CD43 (Siglec-7L), CD52 and CD24 (Siglec-10L) [35–37]. Although Siglec interactions have been studied as novel immune checkpoints, the mechanisms by which they are expressed and function as immunosuppressive checkpoints in the TME are not fully elucidated [38, 39].

Here we assess sialylation across CRC CMS subtype and show that CRC stromal cells express high levels of sialic acid glycans and are enriched for sialylation gene signatures. Sialyltransferases are upregulated in CRC stroma and targeting *ST6GALNAC6* reduces Siglec-10 ligand expression. Stromal-rich CRC is enriched for Siglec-10 and Siglec-10 ligands, and we show that Siglec-10 overexpression in the TME of high fibroblast tumours is associated with poor survival outcome in CRC. We observed that stromal cells induced Siglec-10 expression on primary human macrophages and NK cells. Induction of Siglec-10 is reversed by targeting the Siglec/Sia axis and enhances NK cell cytotoxicity. In a preclinical tumour model, stromal cells induced Siglec-10 on macrophages and pre-targeting of stromal cell sialylation reduced Siglec-G expression by macrophages and increases in CD80-expressing macrophages, as well as granzyme B-expressing and cytotoxic NK cells in tumours. TNF-α-mediated inflammation in tumour-conditioned stromal cells induced Siglec-G expressing macrophages and NK cells in the TME, which were reversed by targeting the Siglec/Sia axis and was associated with lower levels of invasion. Together, this data highlights an integral role for the Siglec receptor/ligand axis in stromal-rich CRC and demonstrates that targeting stromal cell Siglec/Sia interactions modulates innate immune contexture and may represent an innovative strategy to enhance anti-tumour immunity.

## Results

### CRC stromal cells express high levels of sialoglycans

CMS subtype stratification reveals distinctive cellular features of the TME with prognostic implications [40]. CMS4, the mesenchymal subtype, is associated with high stromal infiltration, poor prognosis and limited response to chemotherapy and immunotherapy [3, 40, 41]. We analysed transcriptional profiles of untreated stage II/III colon cancer tumours across CMS molecular subtypes (n=258) (GSE39582) (figure 1A) [42]. Single sample gene set enrichment analysis (ssGSEA) compared the enrichment of pathways related to glycosylation and sialylation (figure 1B). Glycosylation was significantly lower in CMS4 relative to the other subtypes (figure 1C, left). Sialylation-related pathways such as protein sialylation, sialic acid binding, α2,3 sialyltransferase activity, were increased in CMS4 relative to the other subtypes (figure 1C). 2,6 sialyltransferase activity genes were either unchanged or lower in CMS4, (Supplemental figure 1A). We assessed the surface expression of α2-6 and α2-3 linked sialic acid in CRC using immunohistochemistry (figure 1D). Both α2-3 and α2-6-linked sialic acids are expressed in CRC as indicated by SNA-I (α2-6-) and MAL-II (α2-3-) binding (figure 1D, left). SNA-I binding was higher than MAL-II in CRC tissue and MAL-II was significantly higher in the stromal region compared to epithelial region (figure 1D, right), as determined using QuPath (supplemental figure 1B). We assessed α2-3-and α2-6-SA expression in the CMS4 CRC cell lines, SW480, CACO2 and HCT116 and stromal cells using an ex vivo conditioning model with MSCs. MSCs were conditioned with HCT116 tumour cell secretome (TCS) and TNF-α-activated tumour cell secretome (iTCS), (referred to as MSC^TCS^ and MSC^iTCS^) (figure 1E) [20]. Flow cytometric analysis demonstrated that MSCs express higher levels of both α2,3 and α2,6 SA compared to CMS4 like CRC cell lines (figure 1F). MSC^TCS^ express α2,6 SA, which was further enhanced in MSC^iTCS^, while α2,3 SA expression was high and did not change with TCS or iTCS conditioning. Taken together, these data indicate that sialylation is enriched in CMS4 tumours and stromal cells are more highly sialylated.

**Figure 1.**
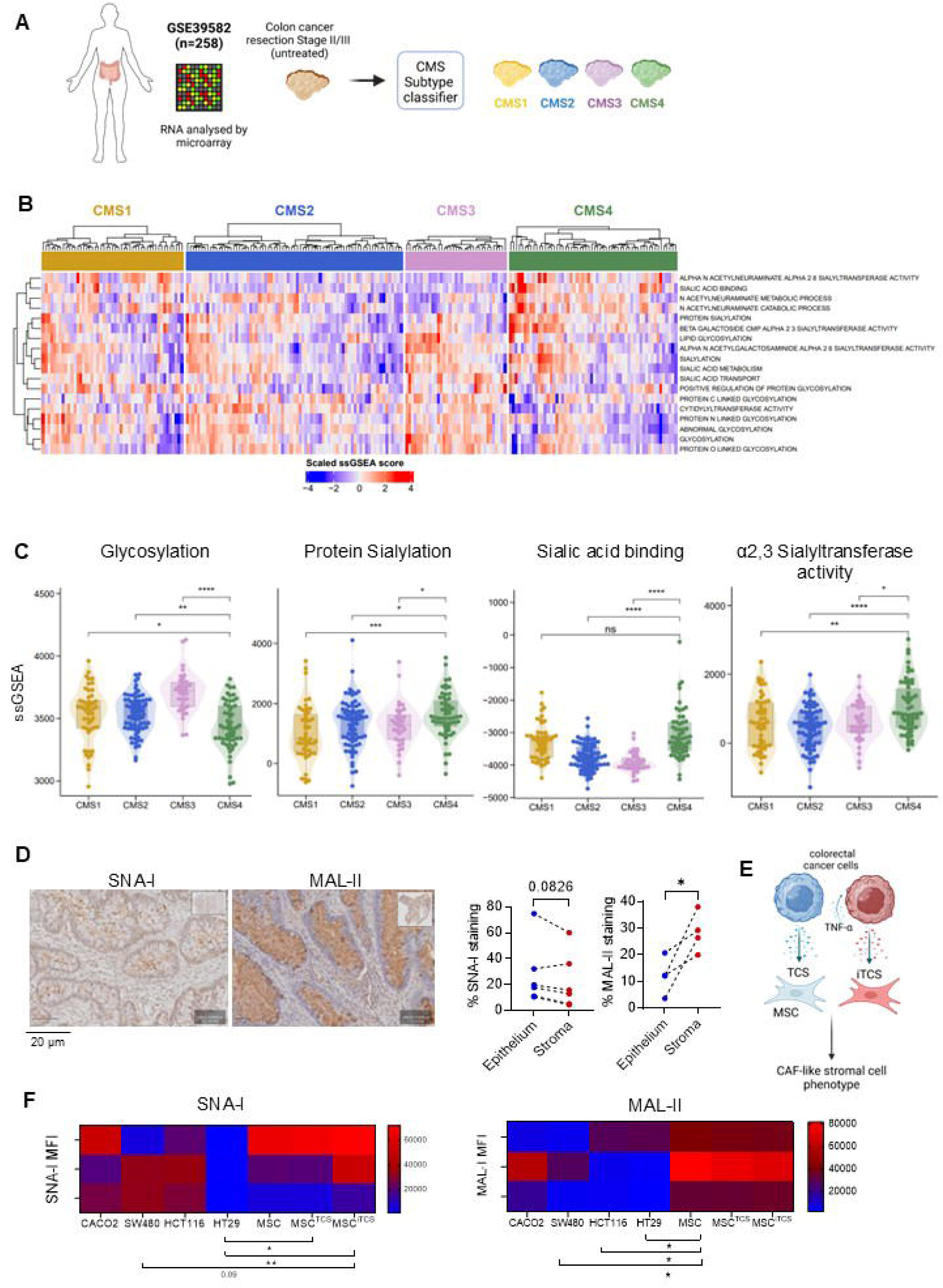
Sialylation is upregulated in CMS4 colorectal cancer. **A** Transcriptional profiles of stage II/III untreated colon cancer samples (GSE39582) were retrieved and CMS classified (n=258; CMS1 = 49, CMS2 = 75, CMS3 = 35, CMS4 = 58). **B** Expression heatmap of transcriptional signatures of glycosylation and sialylation-related genes across CMS1–4 subtypes. **C** Single sample gene set enrichment analysis (ssGSEA) of pathways associated with glycosylation, protein sialylation, sialic acid binding, and α2,3 sialyltransferase activity. **D** Immunohistochemical (IHC) staining images of lectins SNA-I and MAL-II in CRC tissue sections and their quantification in epithelial and stromal regions. **E** Experimental outline of MSC conditioning with tumour cell secretome (TCS) and TNF-α-treated (inflammatory) tumour TCS (iTCS). **F** Heatmaps showing MFI for SNA-I and MAL-II expression in CRC cell lines, MSC^TCS^ and MSC^iTCS^. Data are mean ± SD; *p<0.05, **p<0.01, ***p<0.001, and ****p<0.0001 by Wilcoxon rank-sum test, CMS4 as the reference group (C), paired t-test (D) or one-way ANOVA and Tukey’s *post hoc* test (F).

### Sialyltransferases are upregulated in the stroma of CRC and targeting *ST6GALNAC6* reduces Siglec-10 ligand expression on stromal cells

Sialylation is regulated through sialyltransferase (STs) and neuraminidases (NEUs) activity. We therefore assessed ST and NEU gene expression using ConfoundR [34-36 (https://confoundr.qub.ac.uk/) that enables visualization of gene expression in epithelial and stromal compartments across CRC datasets (GSE35602, GSE39396) (Supplemental figure 2A) [37]. *ST3GAL1-6, ST6GALNAC3, ST6GALNAC5, ST6GALNAC*6 and *ST8SIA1*, were more highly expressed in the stromal compartment compared to the epithelium in most patient samples (GSE53602, figure 2A). NEUs, which cleave sialic acids, were either unchanged between stroma and epithelium (*NEU1,2,3*) or decreased in the stromal compartment (*NEU4*) (figure 2A). Notably, *ST3GAL1, ST3GAL5*, and *ST6GALNAC6* were enriched in fibroblasts and leukocytes (dataset GSE39396; Supplemental figure 2B). We then measured ST expression in primary CRC CAFs isolated from tumour biopsies (Supplemental figure 2C). All ST3 genes, with the exception of *ST3GAL6,* were expressed in CAFs. Only two ST6 genes, *ST6GALNAC4* and *ST6GALNAC6* were expressed (figure 2B). Bulk RNA-seq data from mouse stromal cells showed that *ST6GALNAC6* expression was significantly higher on conditioned MSC^iTCS^ (supplemental figure 2D, E),. Kaplan-Meier plotter analysis (https://kmplot.com/analysis/) [43] shows high *ST6GALNAC6* expression significantly correlated to lower overall survival (OS) in CRC and in CMS4 CRC patients (figure 2C). Furthermore*, ST6GALNAC6* expression was significantly higher in stroma (GSE35602 dataset, figure 2D), and fibroblasts compared to epithelial cells (GSE39396, figure 2E). *ST6GALNAC6* expression was also higher in CMS4 tumours (Supplemental figure 2F) and poor prognosis iCMS3 stromal-rich CRC tumours (figure 2F).

**Figure 2.**
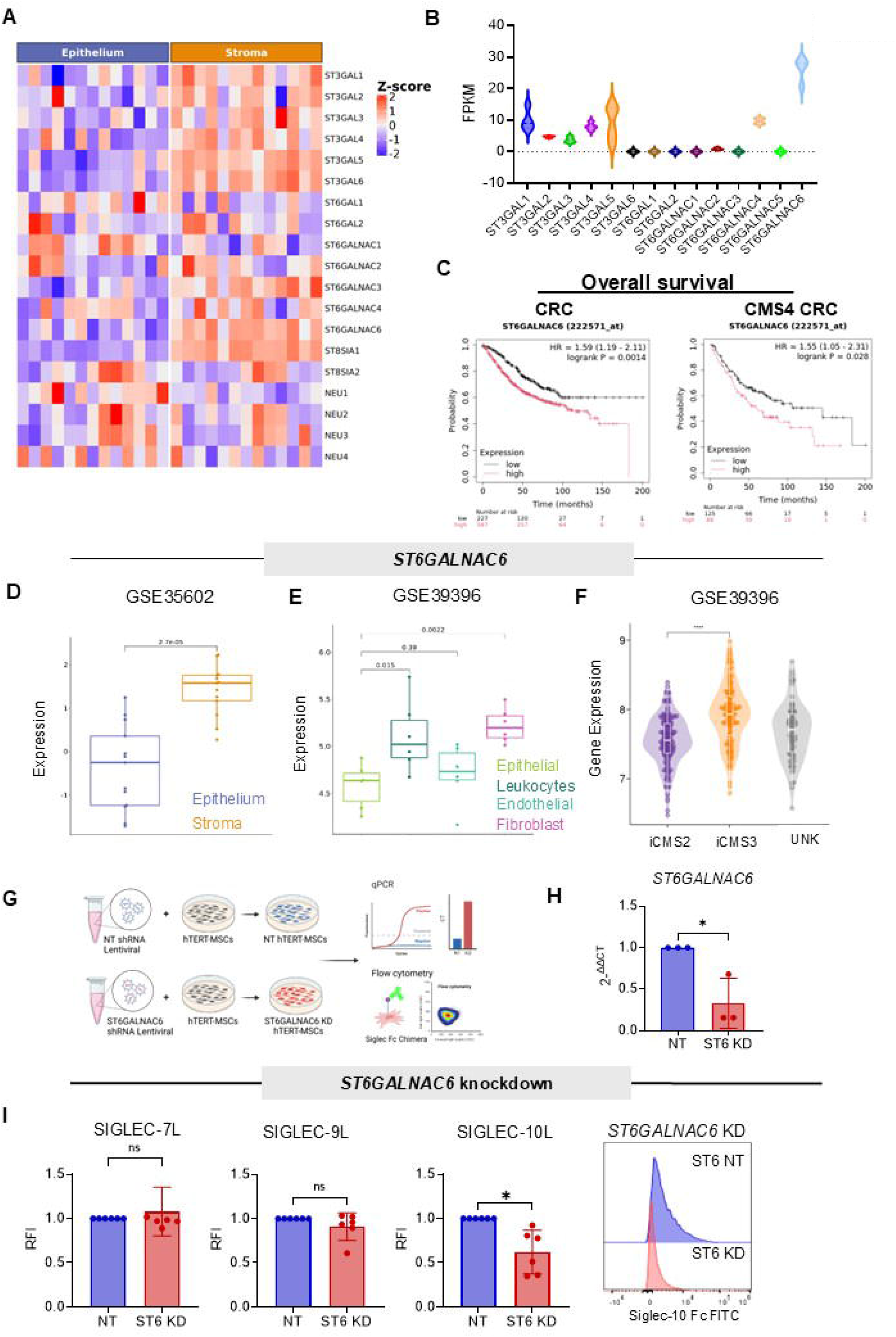
Sialyltranferases are upregulated in colorectal cancer and targeting *ST6GALNAC6* reduces Siglec-10 ligand expression by stromal cells. A Expression heatmap of sialyltransferase and neuraminidase genes in stromal and epithelial compartments in human CRC samples (n = 13) created by ConfoundR (dataset GSE35602). **B** Fragments per kilobase of exon per million mapped fragments (FPKM) values of sialyltransferase genes in human CRC-derived cancer-associated fibroblasts (CAFs) (n = 4), analysed by bulk RNA sequencing. **C** Overall survival (OS) analysis for *ST6GALNAC6* high-and low-expression in CRC (left) and CMS4 CRC samples (right). Kaplan–Meier survival curve showing log rank test *p* value generated using KMPlotter. D-E *ST6GALNAC6* gene expression in different cells of CRC samples created by ConfoundR (datasets GSE35602 and GSE39396). F *ST6GALNAC6* gene expression in iCMS2 and iCMS3 CRC samples (dataset GSE39582). **G** Experimental outline for *ST6GALNAC6* shRNA knockdown in hTERT-MSCs. **H** Relative expression of *ST6GALNAC6* in hTERT-MSCs after shRNA knockdown (KD) compared with non-targeting (NT) shRNA controls (n = 3). **I** Relative fluorescence intensity (RFI) (normalised to NT hTERT-MSCs) of Siglec-7, Siglec-9, and Siglec-10 ligand expression in *ST6GALNAC6* KD hTERT-MSCs (n=6) and representative overlay histograms for Siglec-10 ligand. Data are mean ± SD; *p<0.05, **p<0.01, ***p<0.001, and ****p<0.0001 using one-way ANOVA and Tukey’s *post hoc* test (D-F) or paired t-test (H,I).

We next investigated the role of *ST6GALNAC6* in sialylated ligand production. Using an immortalised stromal cell line, hTERT-MSCs, we generated a stable genetic knockdown (KD) of *ST6GALNAC6* using lentiviral-mediated delivery of shRNA. We confirmed that hTERT-MSCs were highly sialylated; comparable to primary hMSCs, based on α2,3 and α2,6 expression (Supplemental figure 2G). Non-targeting (NT) control was used to assess KD specificity. We confirmed significant reduction of *ST6GALNAC6* gene expression in KD (figure 2H). As Siglec-7,-9 and-10 have been shown to be immunosuppressive in cancer [44], we measured Siglec-7 ligand (Siglec-7L) and Siglec-9 ligand (Siglec-9L) and Siglec-10 ligand (Siglec-10L) expression using Siglec Fc Chimeras. Stromal cells expressed Siglec-7L,-9L and-10L (Supplemental figure 2H) at baseline which was increased by TCS/iTCS (Siglec-7L and-10L). We assessed the effect of *ST6GALNAC6* KD on sialylated ligands and observed no significant differences in Siglec-7L and Siglec-9L expression (figure 2I, left and middle), but we observed a significant reduction in Siglec-10L expression with *ST6GALNAC6* KD compared to NT control (figure 2I, right). Taken together, these data suggest that *ST6GALNAC6* is highly expressed in CRC stromal cells and its expression is involved in the production of Siglec-10L.

### Stromal-rich CRC is enriched for Siglec-10 receptor and ligands and is associated with low relapse-free survival

Siglec receptors are expressed on immune cells and ligand binding triggers inhibitory signals via ITIM and SHP1/2 signalling [45]. In CRC Siglec expression was higher in the stromal compartment than epithelium (Supplemental figure 3A). CMS-classified transcriptional profiles were analysed by a waterfall plot (GSE39582, figure 3A) and boxplot (figure 3B). CMS4 tumours had significantly higher Siglec-10 gene expression compared to CMS2-3 tumours. CMS1 tumours expressed similar levels of Siglec-10 (figure 3B). Siglec-10 expression was significantly higher in high-fibroblast CRC tumours (GSE39582, figure 3C) and positively correlated with fibroblast activation protein (FAP) and podoplanin (PDPN), and negatively correlated with the epithelial marker, EpCAM, (cBioportal, Supplemental figure 3B). High Siglec-10 expression, although not significant, showed a lower probability of relapse-free survival in CMS4-like CRC compared to low Siglec-10 expression (figure 3D). These observations were validated in a second cohort of CRC patients (MTAB 863) (Supplemental figure 3C-G).

**Figure 3.**
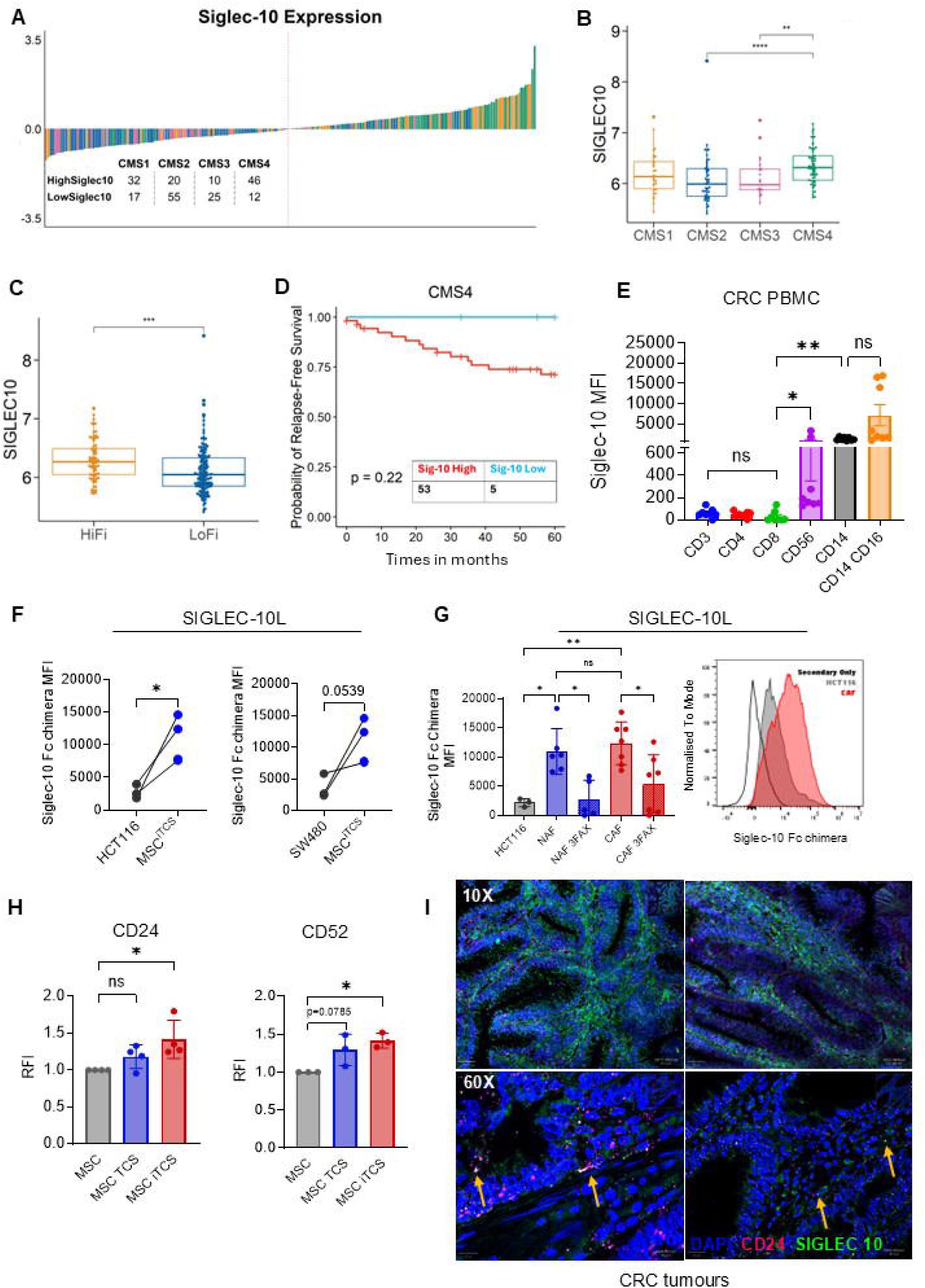
Stromal-rich colorectal cancer enriched for Siglec-10 and Siglec-10 ligand and associated with poor prognosis. **A** Waterfall plot depicting high and low Siglec-10 expression across CMS subtypes (dataset GSE39582). **B-C** Siglec-10 expression levels across CMS subtypes, in high fibroblast (HiFi) and low fibroblast (LoFi) CRC samples (dataset GSE39582). **D** Relapse-free survival analysis for Siglec-10 high-and low-expression in CMS4 tumour samples (dataset GSE39582). Kaplan-Meier survival curves showing log rank test *p* value. **E** Median fluorescence intensity (MFI) of Siglec-10 expression by immune cell subsets in peripheral blood mononuclear cells (PBMCs) from CRC samples (n = 6). **F** MFI of Siglec-10 Fc chimera (Siglec-10 ligand) expression by HCT116 and SW480 CRC cells and MSC^iTCS^ (n = 3). **G** MFI of Siglec-10 Fc chimera (Siglec-10 ligand) expression by CAFs and normal-associated fibroblasts (NAFs) +/-sialyltransferase inhibitor (3FAX) pre-treatment (n=5–6) and overlay histograms showing representative CAF and HCT116 expression. **H** RFI (relative to MSCs) of CD24 and CD52 expression by MSC, MSC^TCS^ and MSC^iTCS^. **I** Confocal microscopy images of multiplex immunofluorescence (M-IF) staining of CD24 (pink) and Siglec-10 (green) in CRC tissue sections (DAPI nuclear staining shown in blue). Data are mean ± SD; *p<0.05, **p<0.01, ***p<0.001, and ****p<0.0001 by Wilcoxon rank-sum test (CMS4 used as the reference group) (B), non-parametric Kruskal-Wallis test (E), ordinary one-way ANOVA and Tukey’s *post hoc* test (F-G).

Next, we measured the expression of Siglec-10 in CRC patient-derived PBMCs, specifically CD3+ T cells (CD4+ and CD8+ T cells) and CD3-cells (CD56+ NK cells and CD14+ and CD16+ monocytes) (gating strategy, supplemental figure 4A,B). Siglec-10 was highly expressed on CD56+ NK cells and CD14+CD16+ monocyte/macrophage subsets (figure 3E). Siglec-10L was increased on MSC^TCS^ and MSC^iTCS^ (Supplemental figure 2H) and was significantly higher on MSC^iTCS^ compared to HCT116 (figure 3F, left) and SW480 (figure 3F, right) epithelial cells.

Sialyltransferases inhibitor (3FAX) 3FaxNeu5Ac, inhibits stromal cell sialylation effectively over a period of 6 days in culture [19]. CAFs expressed higher Siglec-10L compared to HCT116 and the expression was significantly reduced with sialyltransferase inhibitor (3FAX) treatment, confirming sialylation dependent binding (figure 3G). CD24 and CD52, two known ligands for Siglec-10, were also expressed on MSC^iTCS^ (figure 3H) and highly expressed in CMS4 CRC tumours of the FOCUS cohort (Supplemental figure 4C). To assess Siglec-10 localization in CRC tumours, we performed multiplex immunofluorescence (M-IF) staining for Siglec-10 and CD24 which showed Siglec-10 expression was predominantly localized within the stromal compartment of CRC tissue, while CD24 was predominantly in epithelial regions (figure 3I, Supplemental figure 4D). This data shows Siglec-10 receptor and ligand-expressing cells are enriched in the stromal compartment of CRC.

### Stromal cells induce Siglec-10 expression on primary human macrophage and NK cells and inhibit cytotoxicity which can be targeted via the Siglec/Sia axis

Macrophages are one of the most abundant immune cells in the TME of CRC. In CMS4 CRC, macrophages exhibit reduced anti-tumour function [20]. We observed high Siglec-10 expression on macrophages and NK cells and therefore asked whether the Sia/Siglec axis is involved in macrophage-mediated immunosuppression. We observed strong positive correlations between Siglec-10, CD14 and CD16 mRNA expression (cBioportal, CRC TCGA, figure 4A). To understand if the TME impacts expression of Siglec-10 on macrophages, we used an indirect co-culture of primary human macrophages +/-IFN-γ and IL-4 (pro-inflammatory), or IL-4 (anti-inflammatory) with tumour secretome (TCS or iTCS) or MSC^TCS^ or MSC^iTCS^ secretome (figure 4B). We observed no increase of Siglec-10 on naïve, pro, or anti-inflammatory macrophages with tumour cell, HCT116 TCS or iTCS (figure 4C, Supplemental figure 4A). In contrast, Siglec-10 expression was significantly expressed on naïve, pro-inflammatory and anti-inflammatory macrophage in co-culture with MSC^iTCS^ (figure 4D), indicating a stromal-specific increase of Siglec-10 following co-culture with inflammatory CRC stroma. To confirm these effects in primary human NAF and CAF, we set up a direct co-culture with primary macrophages isolated from PBMCs and measured Siglec-10 expression on CD11b+ macrophages (figure 4E). To assess whether this was sialic acid-dependent, NAFs and CAFs were treated with two different sialic acid targeting molecules and strategies, sialyltransferase inhibitor P-3FAX-Neu5Ac (3FAX) (stromal cell pre-treatment) and E610 Bi-sialidase (E610) (Palleons Pharmaceutical) (stromal cell pre-treatment and PBMC co-culture). Both molecules target sialic acid via different mechanisms (figure 4E, bottom). Both 3FAX and E610 effectively reduce stromal cell Siglec ligand expression (Supplemental figure 4E). We observed a significant increase of Siglec-10 expression on CD11b+ macrophages when co-cultured with CAFs, but not with NAFs (figure 4F). Treatment of stromal cell co-cultures with E610 significantly reduced Siglec-10 expression on CD11b+ macrophages (figure 4F, left). But this was not observed with 3FAX pre-treatment of CAFs and NAFs (figure 4F, right). Collectively, these data show that stromal cells selectively induce Siglec-10 receptor expression on primary human macrophages in a sialic acid-dependent manner, which can be reversed by targeting Sia/Siglec axis with E610.

**Figure 4.**
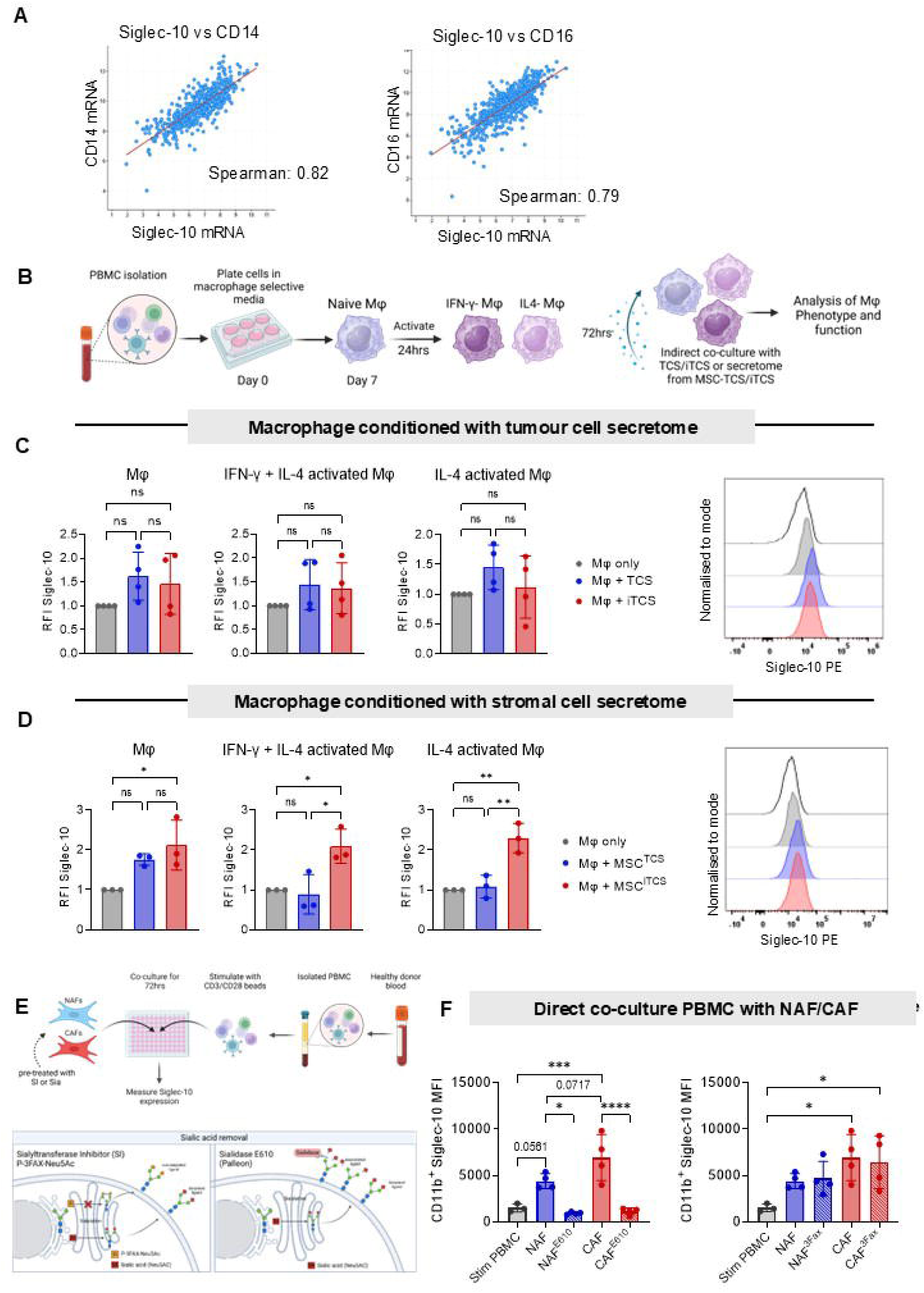
Stromal cells induce Siglec-10 expression on primary human macrophages and are associated with suppressed tumour cell phagocytosis. **A** Dot plots showing correlation between CD14/CD16 and Siglec-10 mRNA expression in a cohort of 394 CRC patient samples from the TCGA database and analysed using cBioPortal (https://www.cbioportal.org). **B** Experimental outline depicting macrophage isolation from PBMCs, cytokine activation and indirect co-culture with cancer or stromal cell secretome. **C** RFI (relative to Mφ alone) of Siglec-10 expression on naïve, IFN-γ and IL-4 activated macrophages after indirect co-culture with HCT116 TCS or iTCS (n=4) with representative overlay histograms. The FMO control is represented by a black line histogram. **D** RFI (relative to Mφ alone) of Siglec-10 expression in naïve, IFN-γ and IL-4 activated macrophages after indirect co-culture with MSC^TCS/iTCS^ secretome (n=3) with representative overlay histogram, The FMO control is represented by a black line histogram. **E** Experimental outline for NAF and CAF direct co-culture with human PBMCs and sialic acid targeting with the sialyltransferase inhibitor (3FAX) and E610-1A sialidase (E610). **F** MFI of Siglec-10 expression by CD11b+ macrophages after co-culture with NAFs and CAFs with or without E610 and 3FAX treatment (n=4). Data are mean ± SD; *p<0.05; **, p<0.01; ***, p<0.001; ****, p<0.0001 using one-way ANOVA and Tukey’s *post hoc* test.

NK cells play an essential role in anti-tumour immunity. They recognize and kill tumour cells without prior sensitization, although NK cell function is often impaired in solid tumours [46] and is suppressed by interactions with stromal cells [47]. NK cells express high levels of Siglec receptors which affects their cytotoxic functions [48]. Siglec-10 was expressed at high levels on CD56+ NK cells from CRC PBMCs (figure 5A). cBioportal analysis showed a positive correlation of Siglec-10 with CD56 mRNA expression in CRC (figure 5B). We co-cultured NAFs and CAFs with stimulated PBMCs (figure 5C) or sorted and expanded human NK cells (figure 5F) and measured Siglec-10 expression on CD56+ NK cells (figure 5D). We observed an induction in Siglec-10 expressing CD56+ NK cells following co-culture with CAFs, which was not affected by pre-treatment of stromal cells with 3FAX, as shown by both frequency and MFI of Siglec-10 on CD56+ NK cells (figure 5E(i)). However, with E610 treatment of stromal and immune cells, we observed a significant reduction in Siglec-10, indicating a sialic acid-dependent mechanism (figure 5E (ii)). Next, we assessed the effects of stromal cells on NK cells isolated and expanded from PBMCs (figure 5F). NK cells were expanded, characterised and co-cultured with NAFs and CAFs. We observed a significant increase in frequency and MFI of Siglec-10 on CD56+ NK cells following co-culture with CAFs compared to NAFs (figure 5G). NK cells were recovered from stromal co-cultures and subsequently co-cultured with HCT116 cells to assess NK cytotoxicity (figure 5H). NK cytotoxicity was unaffected by 3FAX-targeting of NAFs or CAFs (figure 5H, left two graphs). However, NK cytotoxicity was significantly induced by E610 pre-treated CAFs (figure 5H, right). This data shows for the first time that CAFs induce Siglec-10 expression on NK cells and pre-treatment of stroma with sialidase increases NK cell anti-tumour effector cytotoxicity.

**Figure 5:**
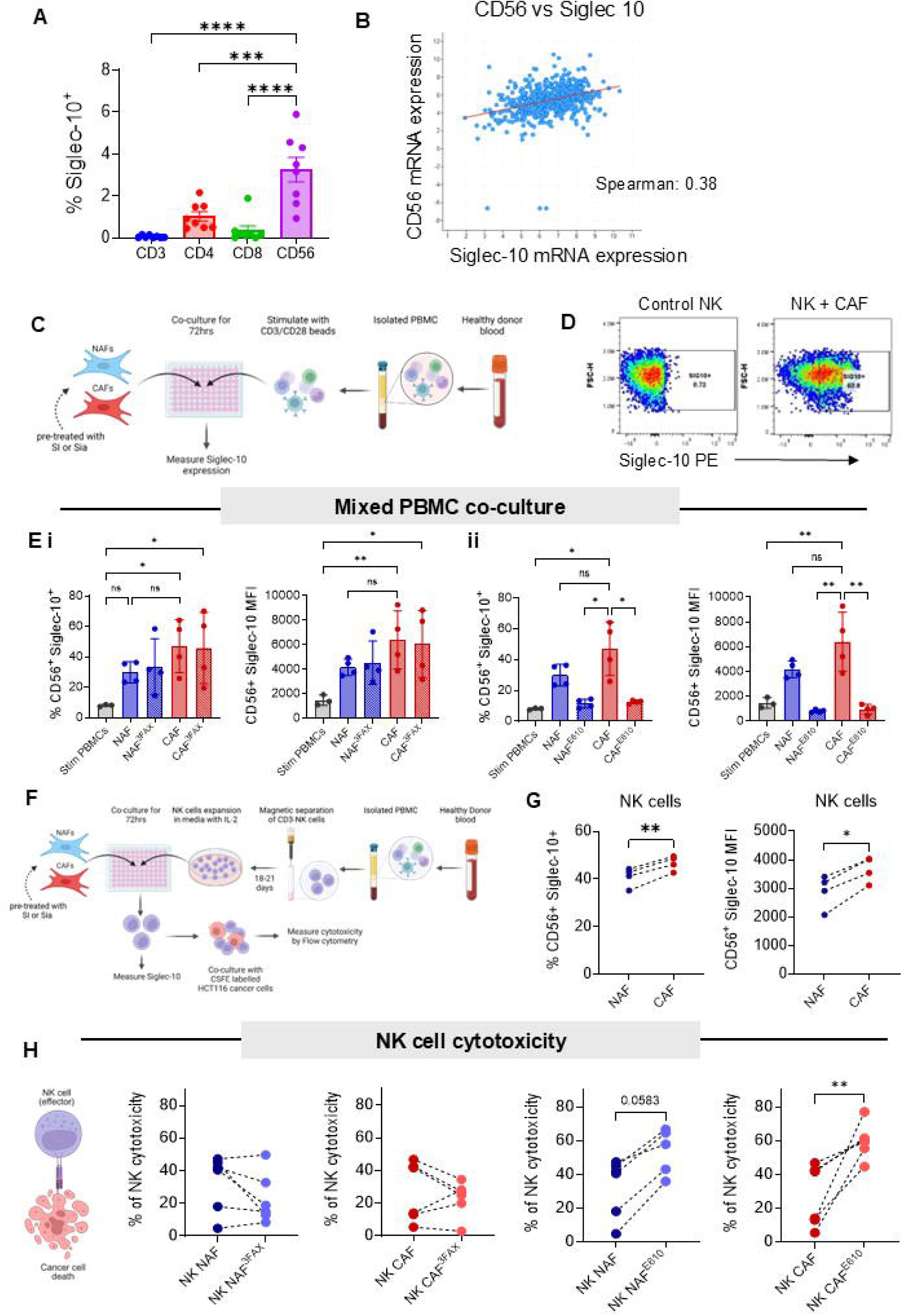
CAFs induce Siglec-10 on primary human NK cells and targeting CAF sialylation increases NK cell anti-tumour function. **A** Frequency (%) of Siglec-10 expression by CD3^+^ T cells, CD4^+^ T cells, CD8^+^ T cells, and CD56^+^ NK cells (n = 8). **B** Dot plot showing the correlation between CD56 (NCAM1) and Siglec-10 mRNA expression in a cohort of 394 CRC patient samples from the TCGA database and analysed using cBioPortal (https://www.cbioportal.org). **C** Experimental outline showing direct co-culture of NAFs or CAFs with stimulated human PBMCs. **D** Flow cytometry dot plots showing Siglec-10 expression by NK cells either cultured alone or following co-culture with CAFs. **E(i)** Frequency (%) (left) and MFI (right) of Siglec-10 expression by CD56+ NK cells after co-culture with NAFs or CAFs with or without pre-treatment with 3FAX (n = 4); **(ii)** Frequency (%) (left) and MFI (right) of Siglec-10 expression by CD56+ NK cells after co-culture with NAFs or CAFs pre-treated and cultured directly +/-E610 (n = 4). **F** Experimental outline showing NK cell isolation from PBMCs and direct co-culture with NAFs or CAFs, with or without 3FAX or E610 pre-treatment, followed by cytotoxicity assay setup. **G** Frequency (%) (left) and MFI (right) of Siglec-10 expression by CD56+ NK cells after direct co-culture with NAFs or CAFs (n=4). **H** Percentage (%) of NK cytotoxicity, measured as the percentage of HCT116 cancer cell killing by NK cells post-co-culture with NAFs or CAFs, with or without 3FAX or E610 pre-treatment (n = 5). Data are mean ± SD; *, p<0.05; **, p<0.01; ***, p<0.001; ****, p<0.0001 by unpaired t-test (G, H) or one-way ANOVA and Tukey’s *post hoc* test (A, E).

### Macrophages *in vivo* are suppressed by CRC stroma which is reversed systemically with sialic acid-targeting

It has been well established that macrophages have prognostic implications in many tumours, including CRC [49]. Using an immunocompetent Balb/c subcutaneous tumour model, we assessed the effects of pre-targeting stromal cell sialylation on innate immune cell phenotypes in tumours, draining lymph nodes (DLN) and spleens at Day 21(figure 6A). To do this, we induced CT26 tumours in the presence or absence of MSC^TCS^ including pre-treatment of MSC^TCS^ with 3FAX or E610. MSC^TCS^ promoted CT26 tumour growth over time, as previously observed [50], and stromal cell de-sialylation led to significantly reduced tumour growth at day 16 (figure 6B). The effect on tumour growth was not fully sustained until day 21, although a significant increase was not seen in MSC^TCS^ ^E610^ at 21 days (figure 6B).

**Figure 6:**
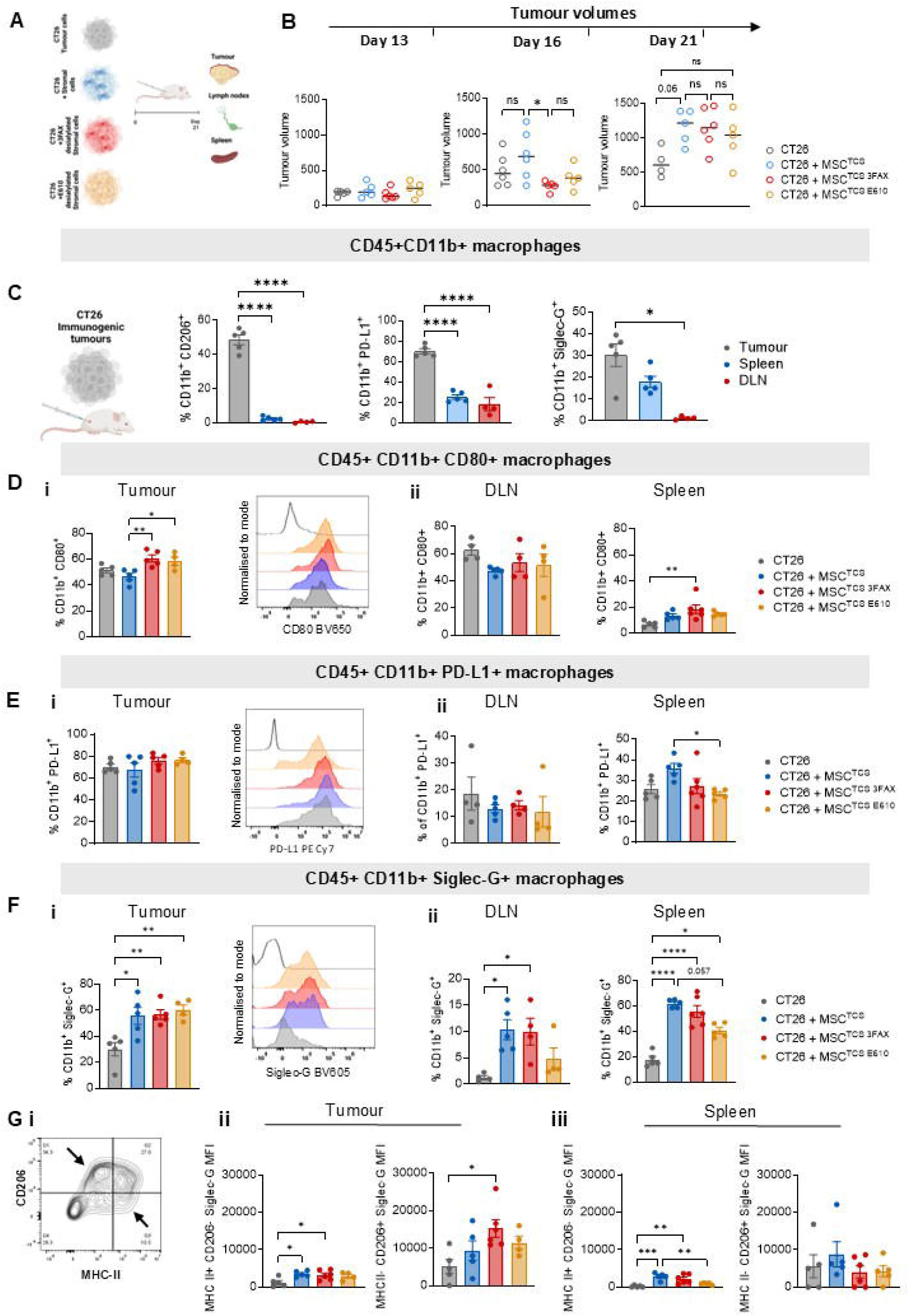
Targeting stromal cell sialylation increases Siglec-G expression by murine macrophages *in vivo*. **A** Experimental outline of murine tumour model. Balb/c mice were injected subcutaneously in the right flank with either CT26 cells alone e, CT26 + MSC^TCS^, CT26 + MSC^TCS^ ^3FAX^ or CT26 + MSC^TCS^ ^E610^ at day 21 post-injection. Tumours, spleens and draining lymph nodes (DLNs) were harvested 21 days post-injection. **B** Tumour volume measured on day 13, day 16 and day 21. **C** Frequency (%) of CD206, PD-L1 and Siglec-G expression by CD45+CD11b+ macrophages in tumours, spleens and DLNs of mice bearing CT26 tumours at day 21 post-injection. **D (i)** Frequency (%) of CD80 expression by CD45+CD11b+ macrophages in murine tumours at day 21 post-injection. Representative overlay histogram (right). **(ii)** Frequency (%) of CD80 expression by CD45+CD11b+ macrophages in DLNs and spleens of tumour-bearing mice at day 21 post-injection. **E (i)** Frequency (%) of PD-L1 expression by CD45+CD11b+ macrophages in murine tumours at day 21 post-injection. Representative overlay histogram shown on the right. **(ii)** Frequency (%) of PD-L1 expression by CD45+CD11b+ macrophages in DLNs and spleens of tumour-bearing mice **F (i)** Frequency (%) of Siglec-G expression by CD45+CD11b+ macrophages in murine tumours. Representative overlay histogram (right). **(ii)** Frequency (%) of Siglec-G expression by CD45+CD11b+ macrophages in DLNs and spleens of tumour-bearing mice **G** (**i)** Flow cytometry contour plot from representative tumour sample showing gating strategy for pro-(MHC II+) and anti-(CD206+) inflammatory macrophage subsets. MFI of Siglec-G expression by MHC-II+CD206-pro-inflammatory and MHC-II-CD206+ anti-inflammatory macrophages in **(ii)** tumours and **(iii)** spleens of tumour-bearing mice at day 21 post-injection. Data are mean ± SD; *, p<0.05; **, p<0.01; ***, p<0.001; ****, p<0.0001 by one-way ANOVA and Tukey’s *post hoc* test. n=4-6 of biological replicates.

We assessed the tumour-associated phenotype of macrophages, specifically CD206, PD-L1 and Siglec-G expression on CD45^+^CD11b^+^ macrophages in the tumour, DLN and spleen by flow cytometry. Macrophages expressing CD206 and PD-L1 are associated with tumour progression, metastasis and immune suppression in CRC [51, 52]. We observed significantly higher frequency of expression of CD206^+^ (figure 6C, Left), PD-L1^+^ (figure 6C, middle) and Siglec-G^+^-expressing (figure 6C, right) CD11b^+^ macrophages in the tumour compared to the secondary lymphoid tissues, indicating that the TME selectively induces a CD206^+^, PD-L1^+^ and Siglec-G^+^ macrophage phenotype. MSC^TCS^ did not significantly change the frequency of CD11b^+^CD80^+^ anti-tumour macrophages in the tumour, however, targeting sialylation increased the frequency of CD11b^+^CD80^+^ anti-tumour macrophages in both SI and sialidase pre-treated stromal cell groups (figure 6Di). This effect appeared to be tumour-specific, as it was not observed in the DLN (figure 6Dii), although an increase in CD11b^+^CD80^+^ macrophages was observed in the spleens of MSC^TCS^ 3FAX-pretretaed group, albeit at much lower frequencies (figure 6Dii). The frequency of CD11b^+^ PDL1^+^ macrophages was higher in tumours compared to DLNs and spleens (figure 6Ei, 6Eii), however, the frequency of CD11b^+^ PDL1^+^ macrophages was unchanged in the tumour and DLN, either by MSC^TCS^ or MSC^TCS^ 3FAX/E610 (figure 6Ei and ii). In the spleen, MSC^TCS^ induced higher frequencies of CD11b^+^PD-L1^+^ macrophages, which was reversed in MSC^TCS^ E610 pre-treated group (figure 6Eii).

We next assessed the expression of the mouse orthologue of human Siglec-10, Siglec-G, on CD11b^+^ macrophages. MSC^TCS^ tumours had significantly higher Siglec-G-expressing CD11b^+^ macrophages, although this was not reversed by targeting stromal cell sialylation (figure 6Fi). We also observed significantly higher levels of Siglec-G-expressing CD11b^+^ macrophages in the DLNs and spleens (figure 6Fii), which was reversed by targeting sialylation in the MSC^TCS^ E610 group specifically (figure 6F ii). These data suggest that targeting stromal sialylation alone is not sufficient to reduce Siglec-G expression on macrophages in the immunosuppressive TME, although Siglec G reduction is evident in the spleen. Finally, we analysed Siglec-G expression on pro and anti-inflammatory macrophage subsets, CD11b^+^MHC-II^+^CD206^-^ and CD11b^+^MHC-II^+^CD206^-^ in tumours (figure 6Gi). Siglec-G expression was lower on pro-inflammatory macrophages than on anti-inflammatory macrophages (figure 6Gii). Siglec-G expression was increased on both pro-and anti-inflammatory macrophage phenotypes in mice co-injected with MSC^TCS^, however, no significant effect was observed by targeting stromal cell sialylation in the tumour (figure 6G ii). Targeting stromal cell sialylation reversed the induction of Siglec-G on pro-inflammatory macrophages in the spleen, but not on anti-inflammatory macrophages (figure 6G iii). Taken together, this data suggests that stromal cells induce an immunosuppressive macrophage phenotype in the CRC TME *in vivo,* characterised by CD206, PD-L1 and Siglec-G expression. Targeting stromal cell sialylation increased frequency of activated macrophages in the tumour and a reduction of immunosuppressive phenotypes in the DLN and spleen indicating a role in regulation of immunosuppression.

### *In vivo* targeting of stromal cell sialylation increases cytotoxic NK cells

In CRC, NK cell infiltration has been identified as a positive prognostic marker in primary and metastatic disease [53]. We therefore assessed NK cell phenotypes in tumour and secondary lymphoid tissues (DLN and spleen) to determine the effect of stromal cell sialylation on NK phenotype and function. NK cells were identified as CD45+CD49b+ cells and NK cell activation marker NKG2D, suppression marker Siglec-G and cytolytic marker granzyme B were also assessed. We observed a higher overall frequency of CD49b+ NK cells, as well as Siglec-G+ and granzyme B+ expressing CD49b+ NK cells in the tumour compared to secondary lymphoid tissues (figure 7A). Notably, NKG2D^+^ activated NK cells were completely absent in the tumour, further supporting the highly immunosuppressive nature of the TME (figure 7A). We next evaluated the effect of stromal cell sialylation on NK cell infiltration. NK cell infiltration in MSC^TCS^ tumours was unaffected, however, targeting sialylation reversed this effect in MSC^TCS^ E610 pretreated groups, significantly increasing NK cell infiltration (figure 7B), which was not observed in MSC^TCS^ 3FAX pretreated tumours (figure 7B). This finding indicates that sialylation of stromal cells plays a role in NK cell recruitment, which has been shown to have prognostic significance in CRC. Despite the increased NK cell infiltration, NKG2D^+^ NK cells were absent in the tumour across all treatment groups, indicating that NK cells within the tumour remain inactivated (figure 7Ci). Interestingly, NKG2D^+^ NK cells were present in the DLNs, with a trend towards higher levels observed in mice co-injected with MSC^TCS^ 3FAX and MSC^TCS^ E610 (figure 7Cii). This same trend was not observed in the spleen (data not shown). This suggests that stromal cell desialylation not only enhances NK cell recruitment but also promotes their local activation in secondary lymphoid tissues, which could be harnessed to optimise therapeutic approaches.

**Figure 7:**
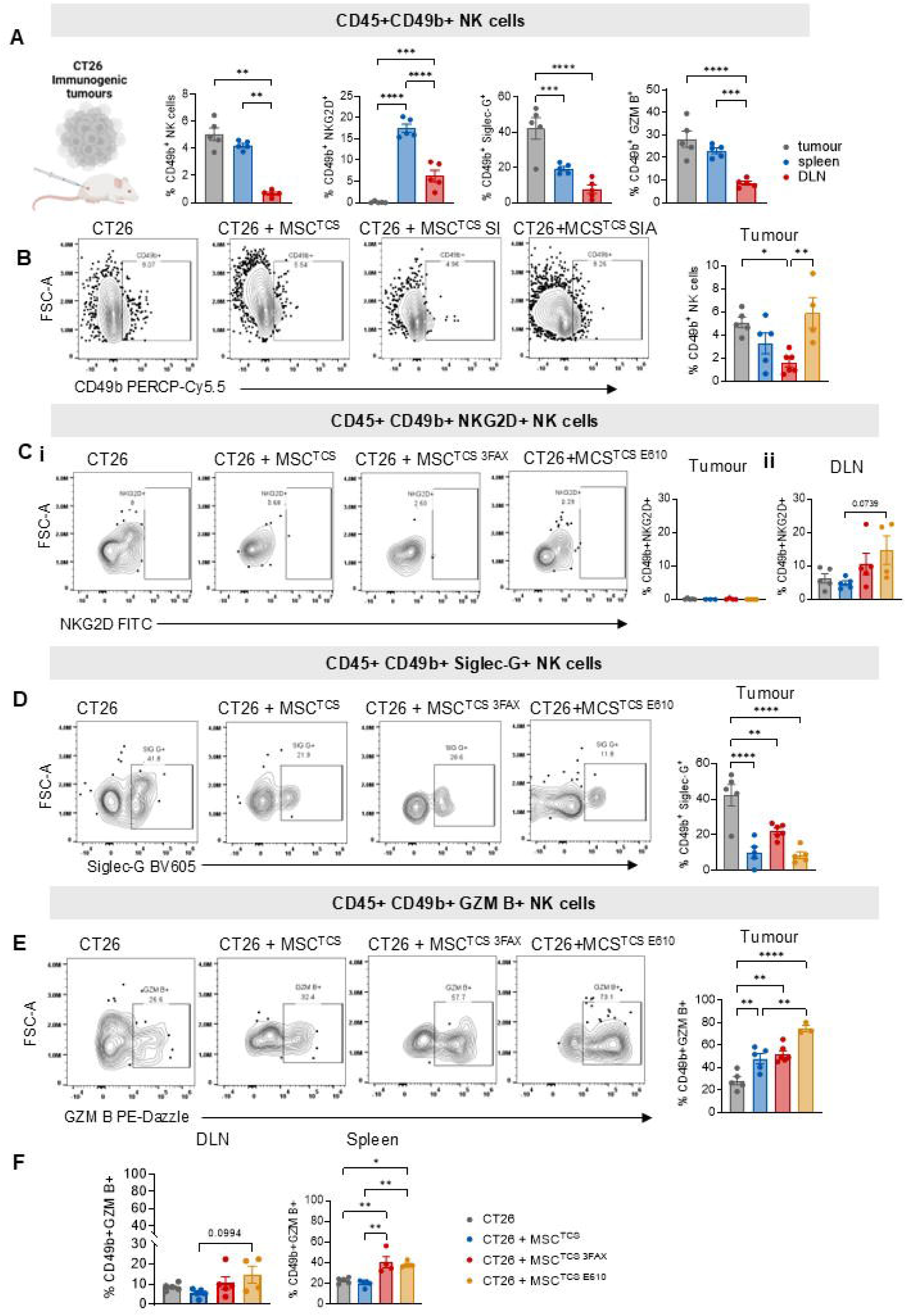
Targeting stromal cell sialylation *in vivo* increases NK cell granzyme B and reduces Siglec-G. **A** Frequency (%) of CD45+CD49b+ NK cells and NKG2D, Siglec-G and granzyme B expression by CD45+CD49b+ NK cells in tumours, spleens and DLNs of mice bearing CT26 tumours. **B** Representative flow cytometry contour plots and quantification of CD49b+ NK cells in murine tumours following injection of CT26 cells alone, CT26 + MSC^TCS^, CT26 + MSC^TCS^ ^3FAX^ or CT26 + MSC^TCS^ ^E610^ at day 21 post-injection. **C (i)** Representative flow cytometry contour plots and quantification of NKG2D expression by CD49b+ NK cells in murine tumours and **(ii)** spleens **D** Representative flow cytometry contour plots and quantification of Siglec-G expression by CD49b+ NK cells. **E** Representative flow cytometry contour plots and quantification of granzyme B expression by CD49b+ NK cells in murine tumours. **F** Frequency (%) of granzyme B expression by CD49b+ NK cells in DLNs and spleens of tumour-bearing mice tumour at day 21 post-injection. Data are mean ± SD; *p<0.05, **p<0.01, ***p<0.001, ****p<0.0001 by one-way ANOVA and Tukey’s *post hoc* test. n=4-6 of biological replicates.

Siglec-G^+^ NK cells were higher in CT26 tumours without stromal cells, reflecting the immunosuppressive TME associated with this model (figure 7D). Frequencies of Siglec-G^+^ NK cells decreased in MSC^TCS^, MSC^TCS^ 3FAX/E610 pre-treated tumours. The activity of NK cells in CRC, especially those expressing granzyme B, correlate with improved survival outcomes [54]. We observed that the frequencies of granzyme B+ NK cells were increased in tumours from mice co-injected with MSC^TCS^ and further significantly increased in tumours from mice treated with MSC^TCS^ ^E610^ but, interestingly, not MSC^TCS^ ^3FAX^ (figure 7E). Similar effects were seen systemically where targeting stromal cell sialylation led to overall higher frequency of granzyme B^+^ NK cells in the spleen (figure 7F), thereby indicating restored cytolytic activity. This was not observed in the DLN, although trend increases are evident in tumours from MSC^TCS^ ^E610^-pre-treated tumours (figure 7F). Our data suggest that stromal cell desialylation may enhance NK cell infiltration and may reprogram them toward an anti-tumour granzyme B-expressing expressing phenotype both locally in the tumour and secondary lymphoid tissues.

### Inflammation alters the immune microenvironment in CRC and induces Siglec-G on macrophages and NK cells which can be reversed by targeting the Siglec/Sialic acid axis

Inflammation enhances stromal cell-mediated immunosuppression and is associated with CRC progression [20, 55]. Analysis of the Biological Hallmark gene sets using GSEA on isolated epithelial and fibroblasts revealed that fibroblasts were significantly and specifically enriched in inflammatory response and TNF-α via NF-κB signalling (GSE39396, ConfoundR) (figure 8A). TNF-α signalling is enhanced in CMS4 CRC (Subtype ExploreR, FOCUS cohort) (figure 8B) [20]. Therefore, to assess the impact of stromal-mediated inflammation on immune cell infiltration and activation, we co-injected CT26 tumours with stromal cells that were treated with iTCS (figure 8C) or MSC^iTCS^ were pre-treated with 3FAX (figure 8C). Tumour cell invasion and NK and macrophage infiltration were assessed at day 13 in tumour, DLN and spleen. We observed a significant increase in the frequency of Siglec-G-expressing macrophages and NK cells in MSC^iTCS^ tumours (figure 8D), which was dependent on stromal cell sialic acid, as it was completely reversed in 3FAX pre-treated stromal cell tumours. Strikingly similar trends were observed in both the DLNs (figure 8E) and spleens (figure 8F) of MSC^iTCS^ and MSC^iTCS^ 3FAX-treated mice. The level of expression of Siglec-G on both CD49b^+^ and CD11b+ cells was significantly higher in the tumour than the DLNs or spleens, indicating a predominant tumour microenvironment effect of immune cell Siglec receptor induction (data not shown). We next assessed Siglec-G expression on CD206+ anti-inflammatory (figure 8G, left) and MHC-II+ pro-inflammatory macrophages (figure 8G, right) and observed an induction of Siglec-G on both subsets of macrophages, which was reversed with 3FAX-pretreatment of stromal cells (figure 8G). Interestingly, 50% of mice co-injected with MSC^iTCS^ showed peritoneal invasion compared to mice injected with CT26 cells alone (figure 8H). This was completely abrogated in mice co-injected with MSC^iTCS^ ^3FAX^. These data show that targeting inflammatory tumour-conditioned stromal cell sialylation inhibits stromal cell-induced metastasis and reduces Siglec-G expression on macrophages and NK cells *in vivo*. We also show that targeting the Siglec/sialic acid axis locally can impact macrophage and NK cell phenotypes outside of the TME.

**Figure 8:**
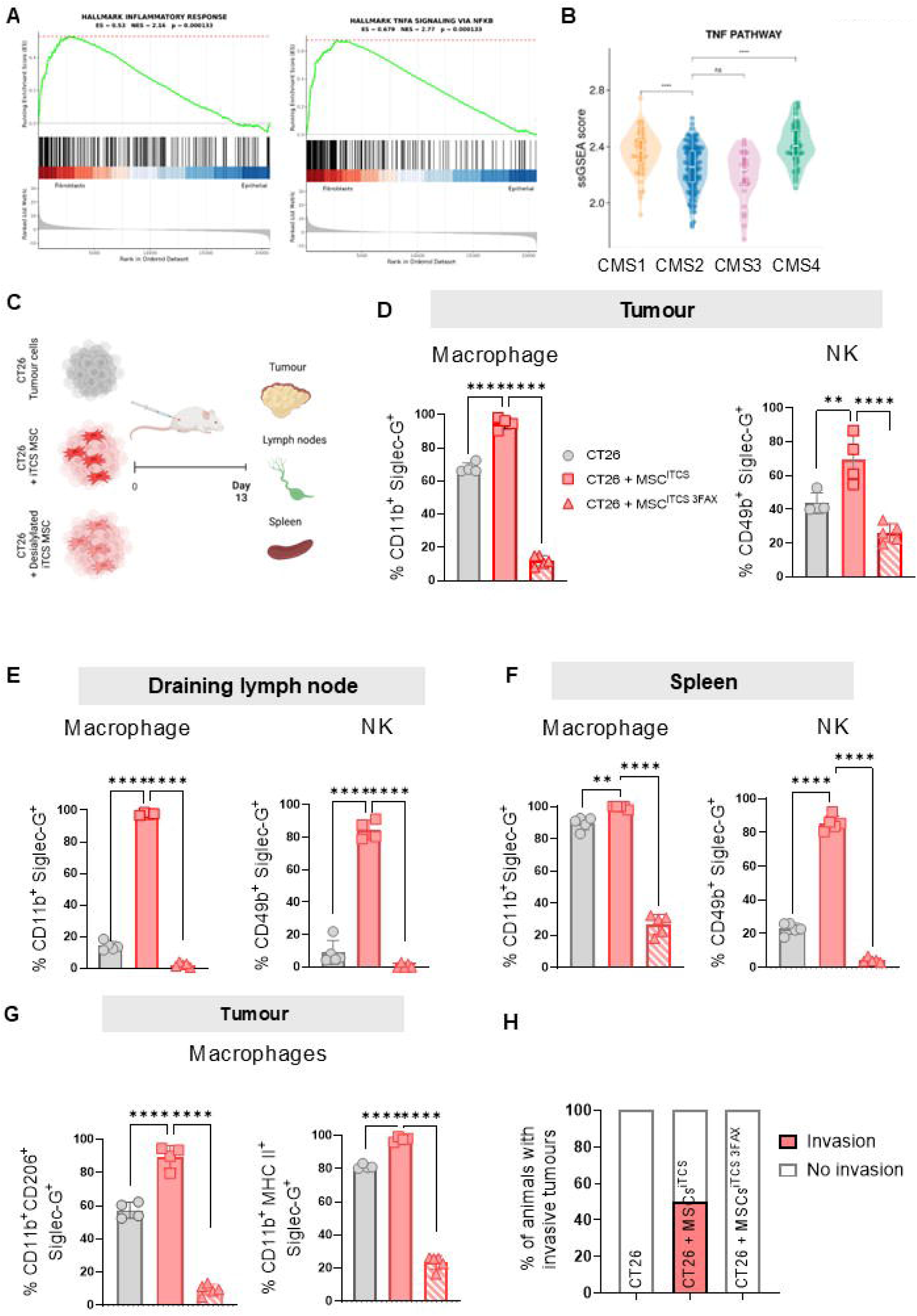
Targeting sialylation of stroma in inflammatory *in vivo* model of CRC reduces Siglec-G expression in tumour-infiltrating macrophages and NK cells. **A** GSEA data depicting hallmark inflammatory responses and hallmark TNF-α signalling via NK-ĸB in CRC tumours comparing transcriptional signatures in fibroblasts versus epithelial cells. **B** Transcriptional signatures of the TNF-α signalling pathway across CMS subtypes 1-4. **C** Experimental outline of murine tumour model. Balb/c mice were injected subcutaneously in the right flank with either CT26 cells alone or co-injected with MSC^iTCS^ or 3FAX pre-treated MSC (MSC^iTCS^ ^3FAX^). Tumours, spleens and DLNs were harvested 13 days post-injection. Frequency (%) of Siglec-G expression by CD11b+ macrophages or CD49b+ NK cells in **(D)** tumours, **(E)** DLNs and **(F)** spleens of tumour-bearing mice. **G** Frequency (%) of Siglec-G expression by CD11b^+^CD206^+^ pro and CD11b^+^MHC-II^+^ anti-inflammatory macrophages in tumours**. H** Percentage (%) of mice with invasive tumours at Day 13. Data are mean ± SD; **p<0.01, ****p<0.0001 using one-way ANOVA and Tukey’s *post hoc* test. n=4-5 of biological replicates.

## Discussion

Sialoglycan-Siglec receptor interactions in the TME play an important role in driving immunosuppression and therapeutic resistance [39]. While aberrant sialylation of cancer cells has been well characterised [38, 56], the sialylation status and function of the tumour-supporting stroma is only beginning to be elucidated in different types of cancer/cancer models. We and others have shown previously that CRC and pancreatic ductal adenocarcinoma (PDAC)-derived CAFs express higher levels of sialoglycans when compared to tumour cells and cancer-adjacent NAFs and these CAF sialoglycans are key modulators of CD8+ T cells and myeloid cells, respectively [19, 24].

Using a combination of co-culture immunoassays, transcriptomic data from CRC patient datasets and an immunocompetent murine model of CRC, we investigated stromal cell sialylation in the context of CMS4 stromal-rich CRC. We assessed the functional impact on stromal cell-mediated modulation of innate immune cells, with a specific focus on macrophages and NK cells. GSEA analysis revealed that gene expression/signatures related to sialic acid binding, protein sialylation and α2,3 sialyltransferase activity were elevated on CMS4-like CRC patient tumours compared to CMS1-3 but, crucially, this same trend was not observed for glycosylation-related gene sets. Given that CMS4-like CRC responds sub-optimally to current therapies and has the worst overall prognosis compared to the other CMS subtypes [5], this information provides insight that hypersialylation may play a role in less-than-optimal response to therapy. In fact, it has been demonstrated previously that sialylation of cell surface glycoproteins can significantly affect the ability of chemotherapy and radiotherapy to eliminate colon cancer cells [57]. While multiple a2,3 and a2,6-specific sialyltransferases are more highly expressed in stroma compared to epithelium in CRC patient samples, *ST6GALNAC6* was the most highly expressed. Furthermore, high expression of *ST6GALNAC6* correlates with significantly poorer survival in both non-classified CRC and CMS4-like CRC. The role of *ST6GALNAC6* in cancer appears to be dichotomous, with some studies reporting decreased expression of this sialyltransferase in CRC [58] and malignant kidney cells [59] compared to normal epithelium, while others show decreased migratory capacity, as an indicator of metastatic ability, of renal carcinoma cells when *ST6GALNAC6* expression is silenced [60]. However, these studies did not assess *ST6GALNAC6* expression in the context of the stroma or stromal-rich tumours. We next sought to determine what effect *ST6GALNAC6* knockdown may have on expression of specific Siglec ligands. Interestingly, we found that shRNA-mediated knockdown of *ST6GALNAC6* significantly decreased expression of Siglec-10 ligands specifically. Therefore, we investigated Siglec-10 receptor expression in the context of stromal-dense CRC. In fibroblast-containing CRC tumours, we show that those with high fibroblast content are significantly higher expressers of Siglec-10 receptor compared to those with a low fibroblastic content. CRC patient tumours with high fibroblastic (Hi-Fi) content, coupled with high Siglec-10 expression, have poorer overall survival compared to Lo-Fi CRC tumours with low Siglec-10 expression. Inflammatory tumour-conditioned MSCs (MSC^iTCS^), show higher expression of Siglec-10 ligand compared to the cancer cells demonstrating that Siglec-10 ligand expression is inducible. In addition, the known Siglec-10 ligands, CD24 and CD52, are also induced and expressed significantly higher on MSC^iTCS^. This prompted us to perform a Siglec-10 receptor screen of different immune cell populations isolated from CRC patient-derived PBMCs. The data revealed that Siglec-10 was significantly more highly expressed on monocytes, macrophages and NK cells.

There is a strong positive correlation between CD14 and Siglec-10 expression, demonstrated by analysis of CRC patient-derived PBMCs and by cBioPortal analysis of CRC patient cohorts. Siglec-10 receptor expression was induced on macrophages cultured in the presence of MSC^iTCS^. The same induction was not observed when macrophages were cultured with HCT116 iTCS alone, indicating that paracrine factors secreted specifically from stromal cells exposed to the inflammatory tumour secretome were responsible for this effect. Two independent strategies were used to target sialylation on fibroblasts to assess what effects desialylation would have on Siglec-10 receptor expression on macrophages. While 3FAX pre-treatment of NAFs and CAFs had no observable effect on macrophage Siglec-10 expression following co-culture, sialic acid cleavage via E610 treatment led to a significant reduction in macrophage Siglec-10 expression. A potential explanation for these contrasting effects may be that 3FAX treatment prevented the trafficking of newly synthesised sialic acid to the fibroblast cell surface, but had no effect on sialic acids already present on the cell surface, while sialidase treatment cleaved sialic acids directly, thereby resulting in reduced Siglec-10 ligand expression on NAFs and CAFs. Further research is required, however, to fully elucidate the mechanism of sialic acid generation in stromal cells. It may also indicate that exposure of immune cells to sialidase is also required for optimal effects.

The second highest expressers of Siglec-10 receptor from immune cell Siglec screening were CD56+ NK cells. cBioPortal analysis revealed a moderate, but significant, correlation between CD56 and Siglec-10 expression in CRC. Similar to our findings on macrophage Siglec-10 expression following co-culture with NAFs and CAFs treated with either 3FAX or E610, we observed the same effects on NK Siglec-10 expression following co-culture in comparable assays. An additional noteworthy observation was that CAFs induced significantly higher frequencies of Siglec-10-expressing NK cells than NAFs following co-culture. Functionally, of the two sialylation-targeting strategies, direct cleavage of surface sialic acids on NAFs and CAFs by E610 yielded the most striking findings with regard to NK cell-mediated cytotoxicity of CRC cells. NK cells co-cultured with E610-treated CAFs were significantly more cytotoxic to CRC cells in subsequent co-culture assays than NK cells that had been co-cultured with untreated CAFs. This effect was not seen when NAFs and CAFs were treated with 3FAX prior to co-culture with NK cells. These data suggest that it may be worthwhile to target ST6GalNac6 specifically on CAFs (e.g. using CRISPR/Cas9-mediated gene knockout) and assess the effects on NK cells, as Kawasaki and colleagues have shown that ST6GalNac6-knockout renal cell carcinoma cells are more susceptible to sialidase-treated NK cell lysis [61]. This may render CAFs more susceptible to killing by NK cells, which may subsequently create a path through the stromal-dense barrier in stromal-rich tumours for other relevant immune cell populations to access the cancer cells and exert their anti-tumour effects.

Using a subcutaneous mouse model of CRC to investigate the effects of co-administration of CRC cells with tumour cell secretome-conditioned stromal cells with or without pre-treatment with 3FAX or E610, we observed lower tumour volumes in mice co-injected with desialylated stromal cells, regardless of the desialylating strategy utilised (i.e. 3FAX or E610), but with a more pronounced effect observed when stromal cells were treated with 3FAX prior to injection. These results show that removal of sialic acids, and potentially specific Siglec ligands, from the stroma, leads to slower tumour growth over time. Similar findings, reported by Scott and colleagues, demonstrated that ST6Gal1 knockdown in prostate cancer cells led to suppression of tumour growth, and by Stanczak et al who showed inhibition of tumour progression in multiple mouse models by targeting tumour cell sialylation through the use of antibody-sialidase conjugates [62, 63]. Assessing novel approaches to targeting sialic acid repeatedly may be more efficacious and give an insight into the kinetics of anti-tumour effects. We profiled specific immune cell subsets within the tumours, as well as key secondary lymphoid tissues (e.g. the draining inguinal lymph nodes (DLNs) and spleens). Siglec-G, the murine orthologue of human Siglec-10 [26], is an inhibitory receptor involved in suppressing anti-tumour immune responses. *Ex vivo* analysis of Siglec-G expression by CD11b+ macrophages within these tissues revealed that Siglec-G was significantly more highly induced in stromal-rich tumours as compared to CT26 tumours alone. A similar effect was seen in both the DLNs and spleen. Interestingly, E610-mediated desialylation of MSC^TCS^ prior to injection resulted in a clear reversal of this induction in secondary lymphoid tissues only. This shows for the first time, to the best of our knowledge, a sialic acid-mediated stromal cell-dependent induction of inhibitory Siglec-G on macrophages in an *in vivo* model of CRC. One possible explanation for why a similar reversal of Siglec-G expression on macrophages was not observed in the stromal-rich tumours in MSC^TCS^ ^E610^ pre-treated mice could be due to the timepoint of this analysis (i.e. day 21 post-injection). Analysis of Siglec-G expression earlier in tumour development would provide further insight into the kinetics of expression and will be investigated in future studies. Our *in vivo* data also provided mechanistic insight into how NK cell phenotype is modulated in stromal-rich CRC tumours and the relative contribution of sialic acid to this modulation. The most striking finding was the almost three-fold increase in granzyme B-expressing CD49b+ NK cells in MSC^TCS^ ^E610^ pretreated tumours. A similar trend was seen in the spleens of tumour-bearing mice. Currently, there is limited information directly linking stromal cell sialylation to the modulation of granzyme B expression by NK cells. Further research is required to understand whether sialylation directly affects NK granzyme B expression or if it influences NK cell activity through interactions with target cells.

Finally the impact of inflammatory conditioning of stromal cells was striking in the promotion of metastatic invasion in CT26 co-injected tumours which was prevented by targeting stromal cell sialylation by 3FAX. These findings highlight that stromal cell sialylation as well as shaping anti-tumour immune phenotypes, may have a role in metastatic invasion of tumour cells. Further research is needed to understand the specific mechanisms involved. Taken together, these findings demonstrate that stromal cells have a strong capacity for innate immune cell modulation in CRC which is regulated, at least in part, by sialylation. We have shown stromal cell-specific induction of Siglec-10 on both macrophages and NK cells and, upon targeting stromal cell sialylation with sialidase, reversal of this effect and a concomitant increase in anti-tumour effector function. Targeting stromal cell sialylation in CRC not only enhances anti-tumour immunity, but also leads to slower tumour growth and prevents metastasis *in vivo*. Targeting stromal cell sialylation in stromal-rich CMS4-like CRC may be the key to reversing immunosuppression and enhancing response to anti-cancer therapies.

<u>Limitations of this study</u>: One limitation of this study is the lack of specificity when targeting stromal cell sialyation, as the two methods utilised (i.e. SI and SIA) inhibit and cleave sialic acids non-discriminately and transiently. Longer-term, vector-mediated knockout of specific sialyltransferases such as *ST6GALNAC6* on stromal cells or the use of monoclonal antibodies targeting Siglec-G or Siglec-10 in tumour xenografts *in vivo* would strengthen these findings.

## Methods

### Human samples

Colorectal cancer-derived NAFs and CAFs were isolated from tumour resections at University Hospital Galway as described [19]. Human PBMCs were isolated from fresh blood samples at University Hospital Galway. PBMCs were isolated by Ficoll-paque (Cytiva Lifesciences) gradient density centrifugation as previously described [19]. Buffy coats containing the PBMCs were collected, resuspended in PBS, and kept on ice for further use.

### Cell lines

Human colorectal cell lines HCT116, HT29, Caco-2 and SW480 are from the American Type Culture Collection (ATCC). HCT116 and HT29 cells were cultured in McCoy’s 5A media (Sigma-Aldrich), SW480 in RPMI-1640 media (Thermo Fisher Scientific) and Caco-2 in DMEM media (Sigma-Aldrich). All media were supplemented with 10% heat inactivated-foetal bovine serum (HI-FBS) (Sigma-Aldrich), 1% L-glutamine and 1% penicillin/streptomycin (Pen/Strep) (both from sigma). Caco-2 were grown in DMEM-low glucose (Sigma-Aldrich) with 20% HI-FBS, 1% L-Glutamine, 1% Pen/Strep and 1% non-essential amino acid (NEAA) (Thermo Fisher Scientific). hTERT stromal cell line was a gift from Dr Maedhbh Brennan. hTERT cells were cultured in DMEM high glucose with sodium bicarbonate, L-glutamine and sodium pyruvate, 10% FBS and 1% Pen Strep (Sigma Aldrich). Human cell master stocks were authenticated by ATCC, confirmed mycoplasma negative, expanded, frozen and used within 20 passages.

### Animal model

Balb/c mice (8–14 weeks, female) were purchased from Envigo Laboratories (Oxon, UK). Animal study was conducted following ethical approval by the Animal Care Research Ethics Committee (ACREC) of University of Galway (ACREC-17-Dec-04) and under individual and project authorisation licences from the Health Products Regulatory Authority (HPRA) of Ireland (AE19125/P077). See supplementary methods for detailed protocol.

### Human mesenchymal stromal cell isolation and tumour cell secretome conditioning

Human bone marrow-derived MSC (hMSCs) were isolated from healthy donors following ethical approval and informed consenr and expanded as previously described [50] and cultured in MEM-α media (Thermo Fisher Scientific) supplemented with 10% HI-FBS, 1% Pen/Strep and 1ng/ml hFGF-2 (PeproTech). The protocol to generate tumour and inflammatory tumour cells secretome (TCS and iTCS, respectively) and stromal cells conditioning was performed according to the method described previously [19].

### Sialyltransferase inhibitor or sialidase treatment

hMSC^TCS/iTCS^ or NAFs/CAFs were treated with 200 µM of the sialyltransferase inhibitor P-3Fax-Neu5ac (3FAX) (Bio-Techne, Abingdon, UK) in two rounds of a 72-hour treatment regimen, as previously described [19] E610-1A sialidase (E610) was provided by Palleon Pharmaceuticals (MA, USA). hMSC^TCS/iTCS^ or NAF/CAF were treated with 100 µg/ml E610 or IgG1 isotype control 24 hours before harvesting for analysis or inclusion in co-culture assays.

### Isolation and polarisation of primary human macrophage

Primary human macrophages were isolated from healthy donor PBMCs as previously described [19, 20]. A total of 8.0×10L PBMCs were plated in 6-well low-adherence plates (Corning) using macrophage complete media (RPMI-1640 supplemented with 1% AB human serum and 1% Pen/Strep). After 2 days, non-adherent cells were washed away, and macrophages were cultured for 7 days with fresh media changes every 2–3 days. For conditioning, 1.5×10L macrophages on day 7 were plated in 24-well low-adherence plates (Corning) and polarised for 24 hours. The media were then replaced with a 50:50 mix of fresh macrophage complete media and TCS or iTCS. Macrophages were conditioned in TCS/iTCS for 2 x 72-hour incubations.

### Isolation and expansion of primary NK cells

Primary human NK cells were isolated from healthy donor PBMCs using a human NK isolation kit and MACS LS column (Miltenyi Biotech) according to the manufacturer’s protocol. Sorted NK cells were expanded in human NK MACS® medium containing 1% AB human serum (ThermoFisher), 1% Pen Strep, and 500 IU/ml human recombinant IL-2 (PeproTech) for 18–21 days, with media changed or split every 2–3 days.

### Transcriptional analysis of CRC datasets

The subset of the stage II/III untreated colon cancer (CC) dataset (GSE39582) [64] (n=258) was previously downloaded and normalized [41]. The samples were classified into the consensus molecular subtypes (CMS) using the CMSclassifier package (1.0.0) (random forest method) [3] ssGSEA scores were generated using the GSVA package (1.38.2) [65]. The gene sets used were retrieved from the Molecular Signatures Database (MSigDB) [66, 67] (accessed November 2022) and imported into R (R 4.0.5). The scores were scaled using the scale() function (default settings) in R with the gene sets as columns. R version 4.3.3 was used for visualisation. The two packages used to generate heatmaps were ComplexHeatmap (2.18) [68] and circlize (0.4.16) [69]. The boxplots were created using ggplot2 (3.5.1) [70], ggbeeswarm (0.7.2) [71] and statistically annotated by ggpubr (0.6.0). Wilcoxon Rank Sum test was used with CMS4 as the reference group. The ConfoundR and Subtype ExploreR web apps were used to generate and download figures. Visualisation for figure 3A-D was created using R version 4.2.2. The previously mentioned cohort, GSE39582 [64], was used. The waterfall plots were created using the ggplot2 (3.5.1). The boxplots were created using ggplot2 (3.5.1) [70], ggbeeswarm (0.7.2) [71] and statistically annotated by ggpubr (0.6.0) [72]. The statistical test used was Wilcoxon Rank Sum test with CMS4 as the reference group. Kaplan Meier plots were created using survminer (0.4.9) (ref 14 – needs added as endnote ref), the cutpoint was obtained using the surv_cutpoint() function and plots were created using the ggsurvplot() function. An additional cohort of stage II primary tumours from CRC patients (n = 215) (E-MTAB-863), previously downloaded and normalized [73] was used to validate bioinformatics analysis in figure 3. These samples were classified into CMS using the CMScaller R package [74].

### Stromal-immune cell direct co-cultures

NAF/CAF and PBMC co-cultures were performed as previously described [19]. Briefly, human PBMCs were resuspended in RPMI-1640 media containing 10% HI-FBS, 1% sodium pyruvate, 1% NEAA, 1% L-glutamine, 1% Pen/Strep and 0.1% β-mercaptoethanol (all Sigma-Aldrich). Cells (1×10L cells/100 μl) were seeded in 96-well round-bottom plates with or without Human T-Activator CD3/CD28 Dynabeads (ThermoFisher Scientific). 3FAX/E610 pre-treated NAFs/CAFs were added to lymphocytes (1×10L cells/100 μl; 1:10 ratio) in complete NAF/CAF medium. E610 was also added to co-culture wells with sialidase pre-treated cells except for the NK cell cytotoxicity assays. After 96 hours, cells were stained with antibodies and analysed using a Cytek Northern Lights Flow Cytometer (Cytek Biosciences). For NK cell co-cultures, 100 μl of NK cell suspension was added to 96-well U-bottom plates, followed by 100 μl of NAF/CAF suspension (1:2.5 NAF/CAF:NK ratio). After 72 hours, NK cells were analysed by flow cytometry or used for cytotoxicity assays.

### NK cytotoxicity assay

NK cells were stained with 50 µL diluted CSFE dye (Thermo Scientific) for 10 mins at 37°C, followed by two washes with NK cell media. Stained NK cells were resuspended in 150 µL complete NK media supplemented with 500 IU/ml human recombinant IL-2 (PeproTech). Simultaneously, HCT116 cells were trypsinized, counted, and resuspended in complete HCT116 media at a concentration of 2.0×10= cells/mL. Then, 50 µL of HCT116 cell suspension was added to U-bottom 96-well plates (Sarstedt), followed by 150 µL of CFSE-stained NK cells to achieve a 2:1 effector-to-target ratio. After 16 hours, cancer cell death was measured by flow cytometry using Sytox Blue staining. Cancer cell death was determined based on CFSE+Sytox+ cells.

### shRNA knockdown of *ST6GALNAC6* in hTERT-MSCs

*ST6GALNAC6* gene knockdown was achieved via lentiviral vector delivery of short hairpin RNA (shRNA). DNA plasmids for lentiviral production of shRNA (GeneCopoeia) were provided as clones encoding shRNA-targeting (KD) and a non-targeting control (NT). The shRNA plasmids were transformed in TOP10F’ *E. coli* cells (ThermoFisher Scientific), expanded in LB broth and the plasmid DNA was isolated using a plasmid Maxiprep kit (Invitrogen). Lentiviral vectors were produced in LentiX293 cells transfected with lentiviral plasmid DNA including shRNA plasmid and three lentiviral packaging plasmids (psPAX2.2, pMD2.G and pRSV-Rev (Addgene)). After 18-24 hours, the culture medium was replaced, and the conditioned medium containing the lentiviral vector was harvested 48 and 72 hours post transfection. Lentiviral supernatant was filtered and used to transduce hTERT immortalized BM-MSCs in T25 flasks. Cells were incubated at 37°C for 72 hours, and GFP expression was measured to assess transduction efficiency. Transduced cells were selected with 50 µg/mL hygromycin (Merck) for 8 days, expanded, and tested for *ST6GALNAC6* knockdown by RT-PCR and flow cytometry.

### Real time PCR (RT-PCR) analysis of ST6GalNAc6 expression

RNA was isolated from *ST6GALNAC6* KD and NT hTERT MSCs using the Invitrogen PureLink RNA Mini Kit (Life Technologies) as per the manufacturer’s instructions. cDNA synthesis was performed using 1 µg of isolated RNA. RT-PCR was performed using TaqMan *ST6GALNAC6* PCR primers (Life Technologies) and was run on the StepOne Plus system (Thermo Fisher Scientific). GAPDH was used as an internal control and relative expression levels were calculated using ΔCT method.

### Flow cytometric analysis of tumour, LNs and spleen immune populations

The protocol for flow cytometric analysis of single cell suspensions isolated from tumour, spleen, draining and non-draining lymph nodes from tumour-bearing mice were described in detail in our previous paper [19]. The detailed protocol and antibody information are provided in the supplementary methods.

### Quantification and Statistical Analysis

All analyses were conducted using Graphpad Prism Version 10 (La Jolla, CA, USA). Experiments were performed in triplicate unless stated otherwise in figure legends. Data was assessed for normal distribution using the Shapiro-Wilk normality test. Datasets containing two groups were analysed by unpaired t-test, Mann-Whitney test or paired t-test. Datasets containing three or more groups were analysed by ordinary one-way ANOVA followed by Tukey’s multiple comparison test, Kruskal-Wallis test followed by Dunn’s multiple comparisons test or two-way ANOVA with Sidak’s multiple comparisons test where appropriate and indicated. P values of <0.05 were considered significant across all statistical tests.

### Key resources table

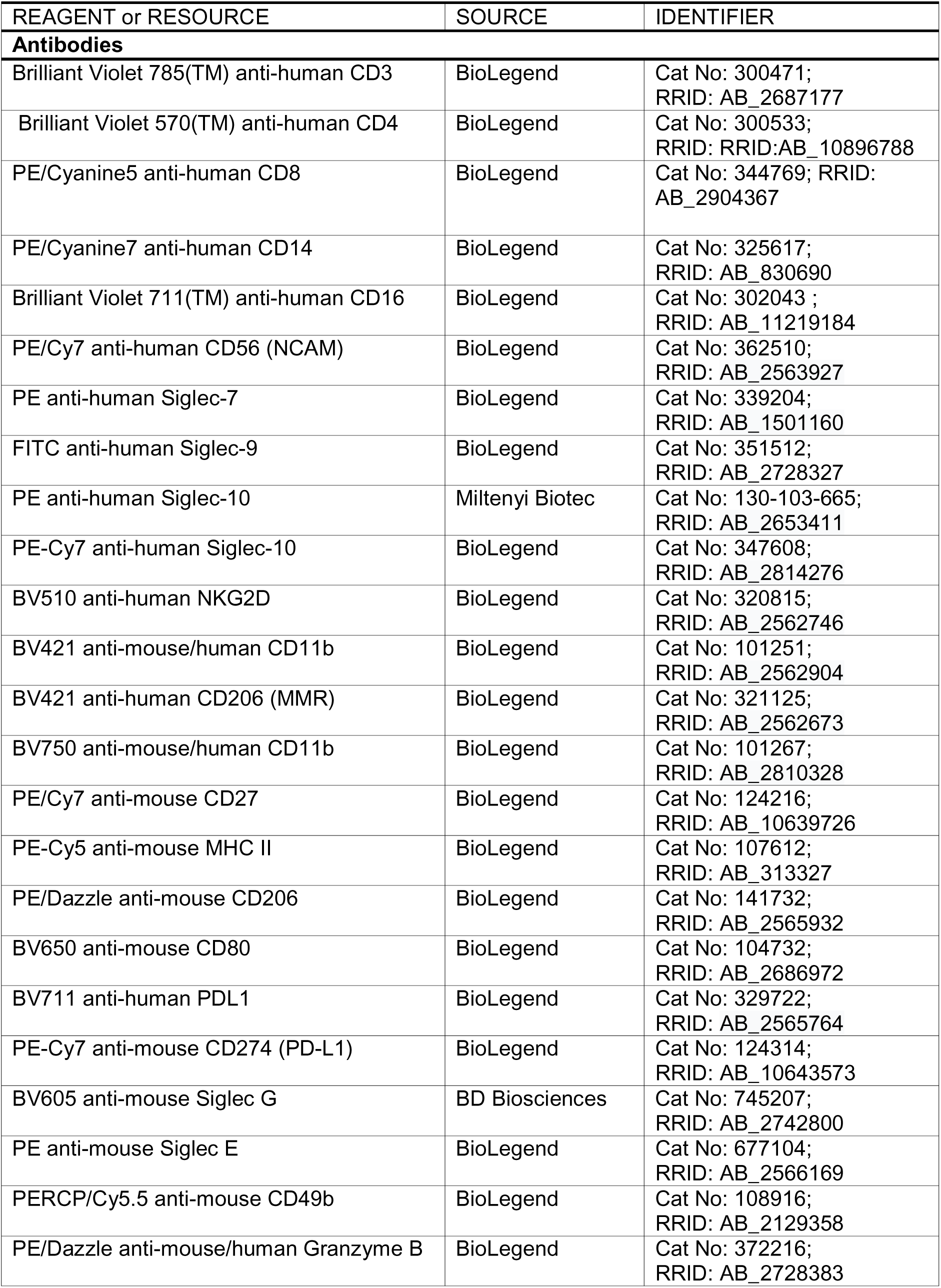
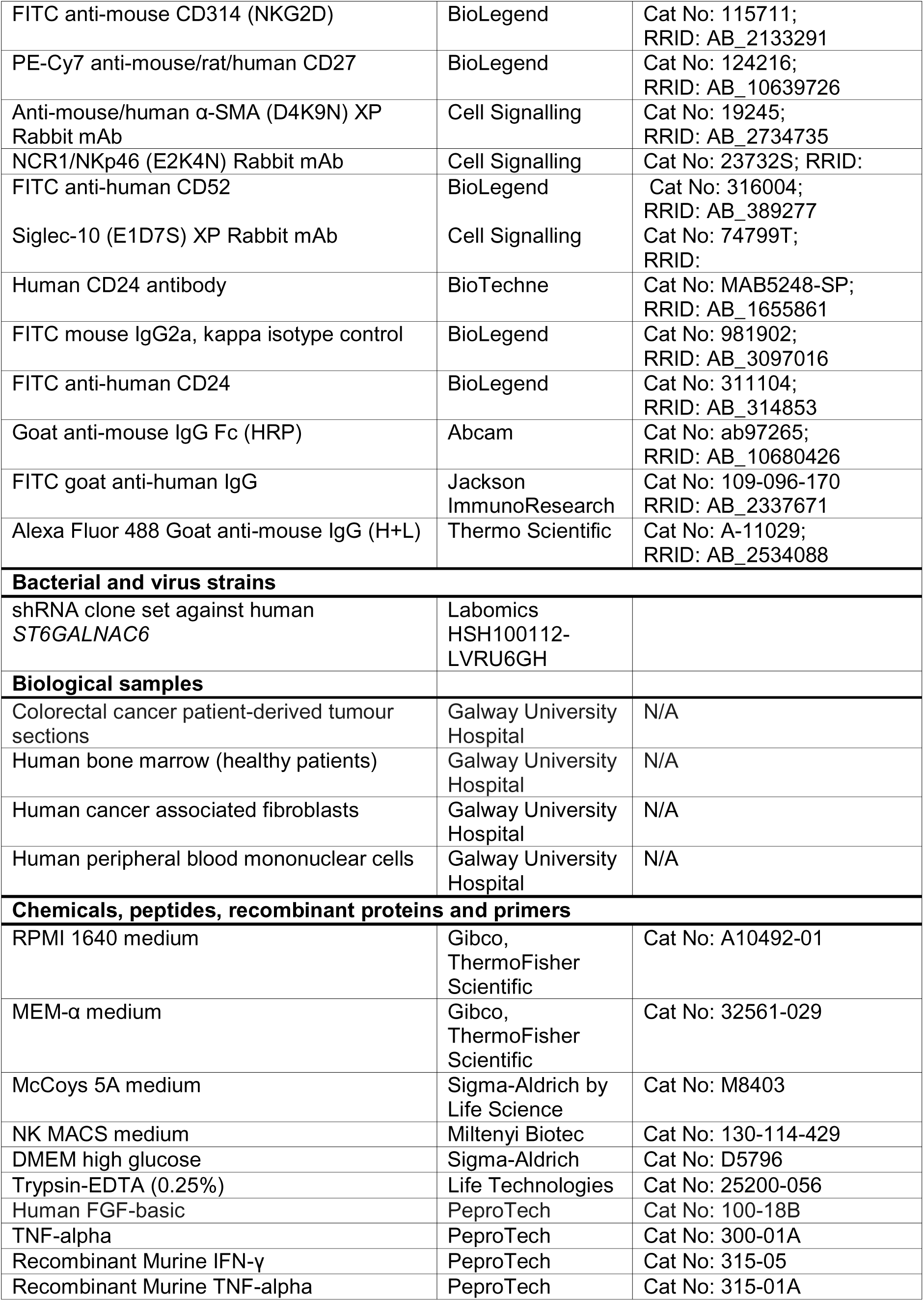
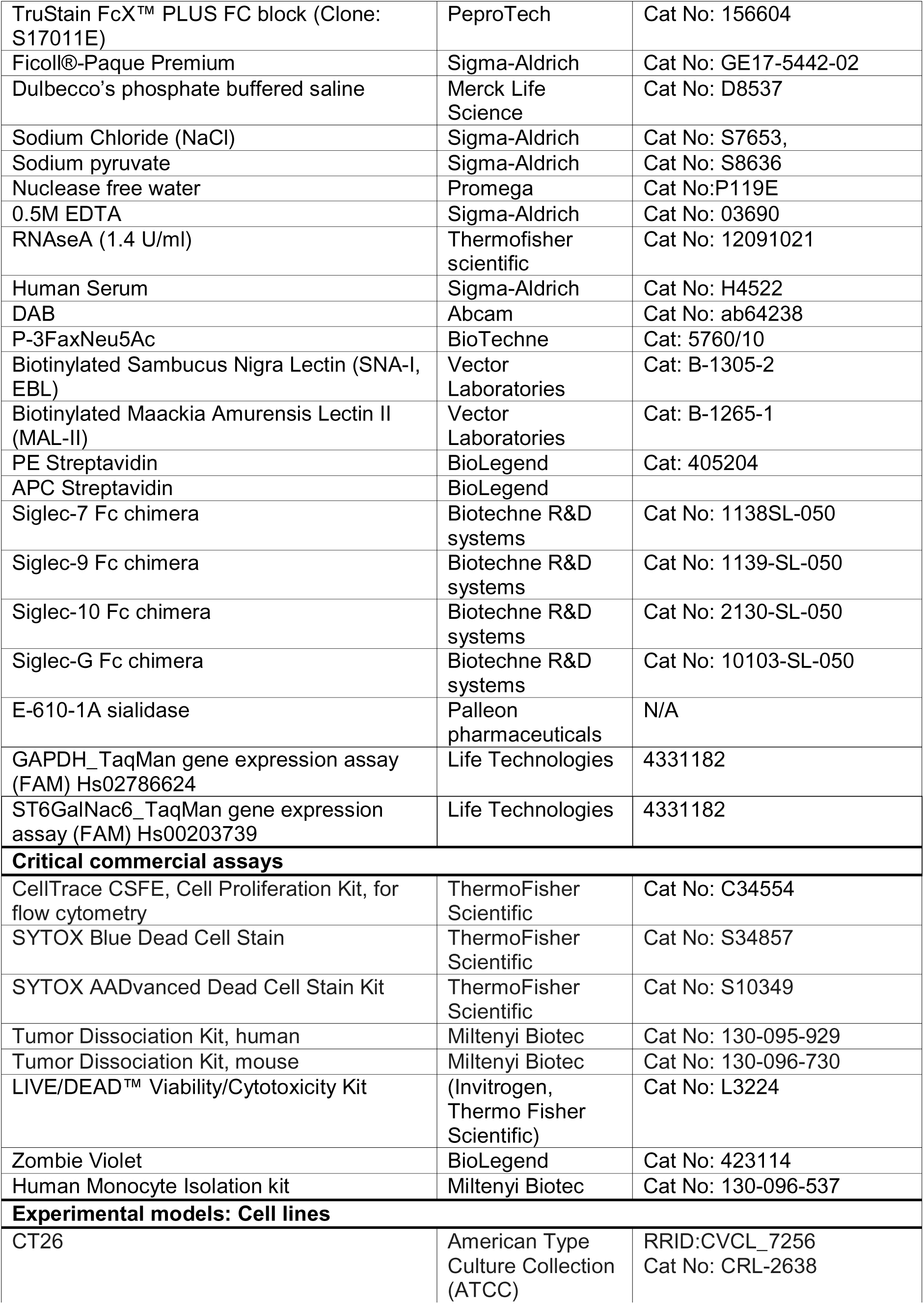
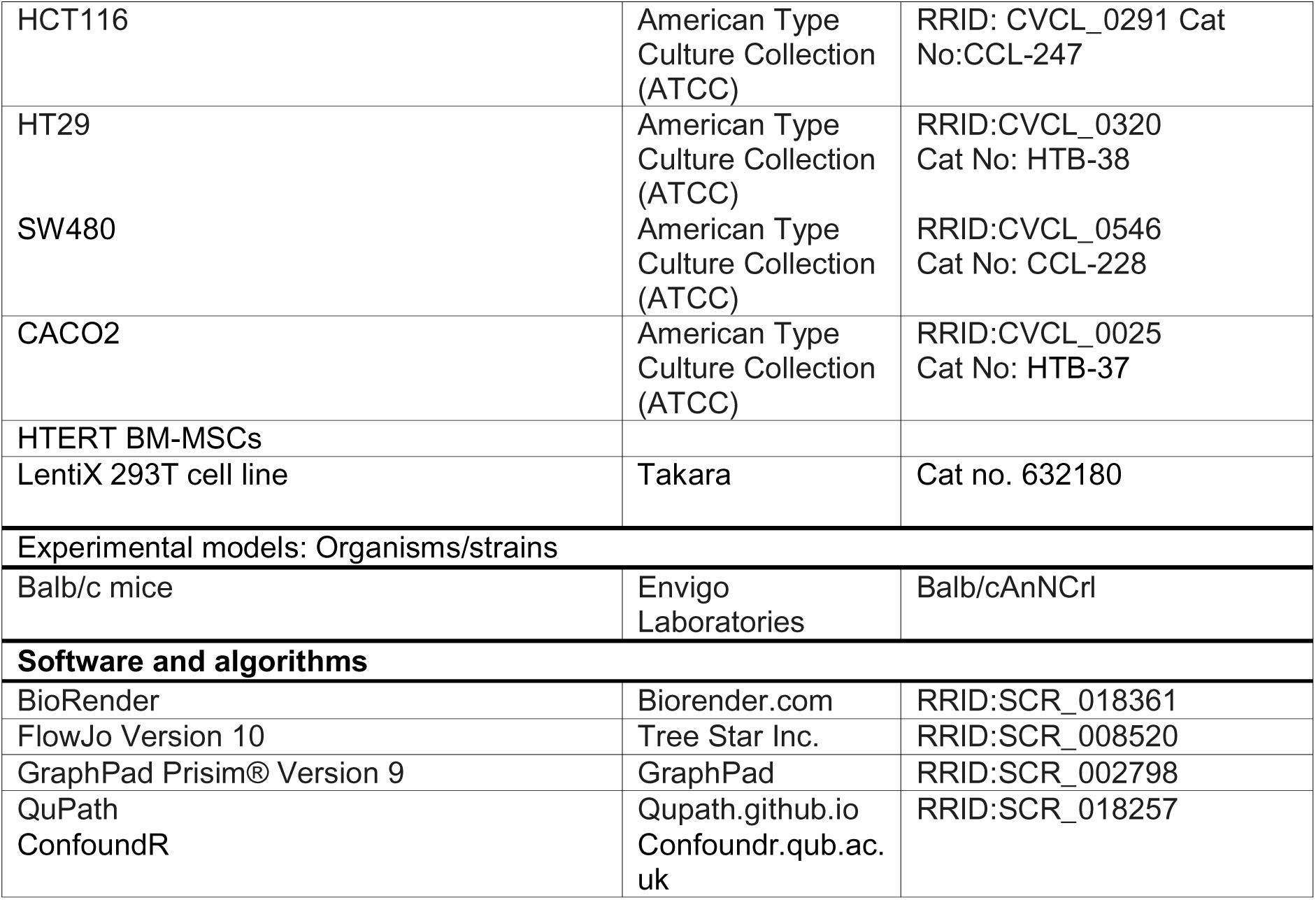

## Declarations

### Ethical approval and consent to participate

Colorectal cancer biopsies and patient-matched blood samples were retrieved from patients undergoing colonic resection at Galway University Hospital under an ethically approved protocol (Clinical Research Ethics Committee, Ref: C.A. 2074 and C.A 3087). All participants gave written informed consent before samples were taken. PBMCs were isolated from the peripheral blood of healthy donors attending a hemochromatosis clinic. Samples were collected under research protocols approved by the Clinical Research Ethics Committee at University Hospital Galway (Ref C.A. 2534). hMSCs were isolated from bone marrow samples following ethical approval (C.A. 02/08-C.A. 429). All participants gave written informed consent before samples were taken.

### Availability of data and material

This paper does not report original code. Any additional information required to reanalyse the data reported in this paper is available from the lead contact upon reasonable request.

## Funding

## Authors’ contributions

A.O’N., N.Z., and H.E performed the majority of experiments and associated data analysis and manuscript writing. S.M.C., C.B., and P.D performed analysis on human CRC datasets and contributed to the interpretation of data and manuscript writing. N.A. L. performed the Siglec screening analysis on immune cells and the associated data analysis. C.O’M and L.H performed the shRNA knockdown experiments. A.W and E.R were involved in the collection of data and processing of PBMCs and tumour biopsies. L.P., L.C. and J.C. provided Sia/Siglec targeting reagents and contributed to the interpretation of data. M.S., A.C., K.C, S.O.H and A.M.H. contributed to study design, planned and performed surgical resections and biopsies and pathological assessment of the samples. L.E., T.R., M.E.O’D., contributed to intial concept and study design, data intrepretation and manuscript review. O.T and A.E.R. conceived the study and contributed to study design, experimental planning, data interpretation, manuscript wiriting.

## Conflicts of interest Statement

L.P., J.C., and L.C. are employees and shareholders of Palleon Pharmaceuticals. M.E.O’D. is a founder of ONK Therapeutics and a member of its Board of Directors and is co-inventor on two related patents (US20210186999A1 and US2017032727899A1). T.R., and A.E.R. are co-inventors on patent US20210186999A1.

## Acknowledgements

The authors acknowledge the facilities and scientific and technical assistance of the Anatomy Imaging and Microscopy Facility at the University of Galway. All flow cytometry experiments were performed in the University of Galway Flow Cytometry Core Facility, which is supported by funds from University of Galway, Science Foundation Ireland, the Irish Government’s Programme for Research in Third Level Institutions, Cycle 5 and the European Regional Development Fund. Technical and consultative support for flow cytometry experiments was provided in the Lambe Institute for Translational Research at the University of Galway by Coralie Mureau and Catherine Loughrey. Graphics and animations created with BioRender, Microsoft PowerPoint and Servier Medical Art. Servier Medical Art by Servier is licensed under a Creative Commons Attribution 3.0 Unported License (https://creativecommons.org/licenses/by/3.0/). The authors wish to thank the Bio-Resources Unit (BRU) technical, veterinary, and administrative staff in the University of Galway for facilitating in vivo studies and for their ongoing assistance, advice, and support in animal procedures, husbandry, care, and welfare. The authors would like to thank Lei Lei and Clodagh O’Neill for the help with *in vivo* study sample preparation. Finally, we wish to acknowledge and thank the patients who, following informed consent, provided samples for this study, for which we are extremely grateful.

## List of Abbreviations

BM-MSC: Bone marrow Mesenchymal stromal cells
CAF: Cancer-associated fibroblast
CMS: Consensus molecular subtype
CRC: Colorectal cancer
DLN: Draining lymph node
ECM: Extracellular matrix
FAP: Fibroblast activation protein
FBS: Foetal Bovine serum
FFPE: Formalin-fixed paraffin embedded
FMO: Fluorescent Minus one
FPKM: Fragments per kilobase of exon per million mapped fragments
GSEA: Gene Set Enrichment Analysis
GZM-B: Granzyme-B
HI-FBS: Heat inactivated FBS
hMSC: human Mesenchymal stromal cell
iTCS: inflammatory tumour cell secretome
MFI: Median fluorescent intensity
MSI: microsatellite instable
MSS: microsatellite stable
mMSC: mouse Mesenchymal stromal cell
NAF: Normal-associated fibroblast
NDLN: Non-draining lymph nodes
RFI: Relative fluorescent intensity
3FAX: 3FaxNeu5Ac Sialyltransferase Inhibitor SIA - Sialidase
Siglec: sialic acid-binding immunoglobulin-like lectin
ssGSEA: single-sample gene set enrichment analysis
TCS: tumour cell secretome
TME: tumour microenvironment

---

**Supplemental Figure 1: Stromal cells in CRC express high levels of sialic acid**

**A** Single-sample gene set enrichment analysis (ssGSEA) of α2,6 sialyltransferase activity in CMS1-4 CRC subtypes from GSE39582 dataset. Wilcoxon rank-sum test, CMS4 used as the reference group. **B** IHC staining images of lectins SNA-I, MAL-II and secondary antibody staining only as negative control for unspecific binding in colorectal cancer tissue sections. Dotted line in black, purple and green showing lumina, epithelium and stroma regions, respectively.

**Supplemental Figure 2: CRC stromal cells express Siglec ligands driven by *ST6GALNAC6* upregulation**

**A** Transcriptional profiles of laser capture microdissected CRC samples (GSE35602 and GSE39396) were analysed using ConfoundR (https://confoundr.qub.ac.uk/). **B** Expression heatmap of sialyltransferase and neuraminidase genes in epithelial cells, leukocytes, endothelial cells and fibroblasts in human CRC samples (n = 13) analysed by ConfoundR (GSE35602). **C** Experimental outline showing isolation of CAFs from CRC biopsies. **D** Experimental outline showing conditioning of murine MSCs with TCS and iTCS from CT26 cancer cells. **E** FPKM values of *ST6GALNAC6* gene expression in MSC and MSC^iTCS^ from bulk RNA-sequencing. **F** *ST6GALNAC6* gene expression in CMS1-4 tumour subtypes from GSE39582 dataset. **G** Overlay histograms of SNA-I and MAL-II expression by primary human MSCs, intestinal MSCs (iMSC#3) and hTERT immortalized MSCs (hTERT-MSC). **H** MFI of Siglec-7,-9 and-10 ligand expression by MSCs, MSC^TCS^ and MSC^iTCS^ analysed via flow cytometry. Data are mean ± SD; *p<0.05, **p<0.01, ***p<0.001 by paired t-test (E), one-way ANOVA and Tukey’s *post hoc* test (H). n=5.

**Supplemental Figure 3: Siglec-10 and Siglec-10 ligands are associated with CRC stroma**

**A** Expression heatmap of Siglec receptor genes in the epithelium and stromal compartments of human CRC samples (n = 13) created by ConfoundR (GSE35602),left). **B.** Bar graphs showing statistical comparisons of Siglec 7,9 and 10 (GSE35602), right). **C** Dot plots showing correlation between FAP/PDPN/EpCAM and Siglec-10 mRNA expression in a cohort of 394 CRC patient samples from the TCGA database and analysed using cBioPortal (https://www.cbioportal.org). **D-F** Waterfall plot and boxplots showing Siglec-10 expression across CMS1-4 and in high fibroblast (HiFi) and low fibroblast samples in the validation cohort MTAB 863. **G** Kaplan–Meier curve showing relapse free-survival in Siglec-10 high and low samples in the validation cohort MTAB 863.

**Supplemental Figure 4: Siglec-10 and Siglec-10 ligands are associated with CRC stroma**

**A** Experimental outline of Siglec receptor analysis in PBMCs isolated from peripheral blood of CRC patients. **B** Flow cytometry gating strategy used to analysed single, live, and CD3-(CD56+ NK and CD14+ and CD16+ monocyte) and CD3+ cells (CD4+ and CD8+ T cells). **C** CD52 gene expression in CMS1-4 CRC samples in the FOCUS cohort. **D** Multiplex immunofluorescence images of CD24 (pink) and Siglec-10 (green) staining in CRC tissue sections and in normal colon tissue section (blue indicate DAPI nuclear staining). **E** Bar graphs showing Flow Cytometry data of Siglec 7L expression on MSC +/-conditioning and treated +/-3FAX and E610 (left) Overlay histograms showing overall level of expression and efficacy of reduction by 3FAX and E610 (n=3)

**Supplemental Figure 5: Siglec-10 expression on macrophages is not cancer cell dependent**

**A** RFI (relative to M0 macrophage) of Siglec-10 expression in naïve (M0), IFN-γ and IL-4 stimulated (M1) and IL-4 stimulated (M2) primary human macrophages. **B** Flow cytometry gating strategy used to analyse single, live and Siglec-10 expression by CD11b+ macrophages following co-culture with MSC^TCS^ and MSC^iTCS^.

## Notes

### Competing Interest Statement

L.P., J.C., and L.C. are employees and shareholders of Palleon Pharmaceuticals. M.E.OD. is a founder of ONK Therapeutics and a member of its Board of Directors and is co-inventor on two related patents (US20210186999A1 and US2017032727899A1). T.R., and A.E.R. are co-inventors on patent US20210186999A1.

