## Supplementary methods for "Stromal cells modulate innate immune cell phenotype and function in colorectal cancer via the Sialic acid/Siglec axis"

**Subcutaneous tumour model**

Balb/c mice (aged 8–14 weeks, female) were purchased from Charles River UK (Kent, UK). Ethical approval for this research was granted by the Animal Care Research Ethics Committee (ACREC) committee of the University of Galway (ACREC-17-Dec-04). Experiments were conducted under individual and project authorisation licences from the Health Products Regulatory Authority (HPRA) of Ireland (AE19125/P077). Female mice were acclimatised for at least 7 days prior to tumour induction and randomised to specific groups on Day 0. All mice were housed in individually ventilated cages under specific pathogen-free conditions, fed ad lib on a standard chow diet, and cared for under the standing operating procedures in the BRU Animal Facility at the University of Galway, Ireland. Sample size calculations for animal numbers per group were performed prior to experiments.

For tumour experiments, 5x10^5^ CT26 tumour cells were inoculated subcutaneously (s.c.) in the flank of mice under 2-3% isoflurane ±1.5x10^5^ MSC^TCS^ or MSC^iTCS^ with or without pre-treatment with sialyltransferase inhibitor (3FAX) (Biotechne) or sialidase E610-1A (Palleon Pharmaceuticals) in a total volume of 100 µl. Tumour growth was monitored daily, and size was measured 3x per week using a digital callipers. Tumour volume was calculated according to the rational ellipse formula: (M^1^2^ x M^2^ x π/6). Mice with a CT26+/- MSC^iTCS^ tumour were euthanised at day 13, while mice with a CT26 tumour +/- MSC^TCS^ +/- 3FAX/E610 pre-treatment were euthanised at 21. At endpoint, tumours, draining lymph nodes (DLN), non-draining lymph nodes (NDLN) and spleens were harvested. Mice were euthanised at endpoint or if tumours showed signs of ulceration or if clinical endpoint was reached. In the case of ulcer formation, mice were excluded from study analysis.

### **Immunohistochemistry (IHC) and Immunofluorescence (IF)**

Formalin-fixed paraffin embedded (FFPE) human tumour tissue was kindly provided by the Histopathology Department at Galway University Hospital. Slides were immersed in xylene and decreasing percentage of ethanol (100%, 70%, 30%) followed by immersion in water. Antigen retrieval was performed in a pressure cooker at 95°C for 20 mins in sodium citrate buffer, pH 6.0. Slides were left to cool then endogenous peroxidase activity was blocked using 3% hydrogen peroxide (H_2_O_2_) for 5 mins. Protein blocking was performed using Tris-Buffered saline (TBS) with 3% goat serum for 45 mins at room temperature (RT). During blocking, HYDRA reagents (Palleon pharmaceuticals) were prepared at optimal concentrations in TBS with 3% goat serum. HYDRA-3, -7 and -9 were added to each slide and slides were incubated at 4°C overnight. The next day, slides were washed for 15 mins in TBS-T to remove unbound antibody. For IHC, the slides were stained with secondary antibody goat anti-mouse HRP for 1 hr at RT, washed and then stained with DAB chromogen for 5 mins, washed and counter-stained with hematoxylin for 1 min. Then slides were rinsed in water and dehydrated using an ethanol series and xylene. For IF staining, after primary Ab incubation, the slides were stained with secondary antibody goat anti-mouse APC for 1.5 hrs at RT in the dark, washed and stained with DAPI nuclear counterstain for 5 mins. All slides were mounted with DPX and left at 4°C overnight before imaging using an Olympus slide scanner VS120.

### **Flow cytometric analysis of immune populations in tumours, DLNs, NDLNs and spleens**

To analyse tumour immune cell infiltrates, tumours were dissociated using a mouse Tumour Dissociation Kit (Miltenyi Biotec) according to the manufacturer’s protocol with modifications, and incubated for 2 hrs at 37ºC with inverting 10 times every 30 mins. Single-cell suspensions of dissociated tumours, draining and non-draining lymph nodes and spleens were prepared by gentle mashing of the organs through 40 µm cell strainers (ThermoFisher Scientific) that were placed on top of 50 mL tubes (Sarstedt) containing 5 ml of DPBS. The single-cell suspensions were then centrifuged at 400 x g for 5 mins. Lymphocytes were washed in DPBS and counted using a hemocytometer. Splenocytes and tumour-infiltrating lymphocytes were re-suspended in ACK lysis buffer (Sigma-Aldrich) and incubated on ice for 5 mins. The reactions were stopped by adding 10x complete medium consisting of RPMI 1640 (ThermoFisher Scientific) supplemented with 10% heat-inactivated FBS, 1% sodium pyruvate (1mmol/L), 1% non-essential amino acids (0.1mmol/L), 1% L-glutamine (2mmol/L), 1% penicillin (100U/ml)/streptomycin (100mg/ml), and 0.01% β-mercaptoethanol (55mmol/L) (all from Sigma-Aldrich). Cells were centrifuged at 400 x g for 5 min, washed, and re-suspended in DPBS and counted. For flow cytometric analysis, 1.0x 10^5^ cells per sample (and appropriate fluorescence minus one (FMO) controls) were stained with the following anti-mouse antibodies diluted in fluorescence-activated cell sorting (FACS) buffer (DPBS supplemented with 2% FBS and 0.05% sodium azide): CD45.2-BV510, CD49b-PerCP/Cy5.5, CD11b-BV750, CD27-PE-Cy7, NKG2D-FITC, CD206-PE-DAZZLE, MHC-II-Pe-Cy5, PD-L1-PE-Cy7, CD80-BV650 and Siglec-G-BV605 (all Biolegend) (see key resources table for antibody catalog numbers) and with the cell viability dyes SYTOX Blue or Zombie Violet (ThermoFisher Scientific). For granzyme B staining, cells were washed twice with DPBS after surface staining and fixed (2% paraformaldehyde (Sigma-Aldrich) for 10 minutes at room temperature (RT) for intracellular staining. Cells were then incubated with anti-mouse granzyme B antibody diluted in permeabilization buffer (1% BSA/PBS with 0.5% saponin) according to the manufacturer’s instructions. Samples were incubated at 4°C for 10 mins and washed twice in FACS buffer. Samples were analyzed using a Cytek Northern Lights Flow Cytometer. Flow cytometry data was analysed using FlowJo analysis software version 10.
