## Supplementary figures and images for "Stromal cells modulate innate immune cell phenotype and function in colorectal cancer via the Sialic acid/Siglec axis"

### Supplementary Figure 2

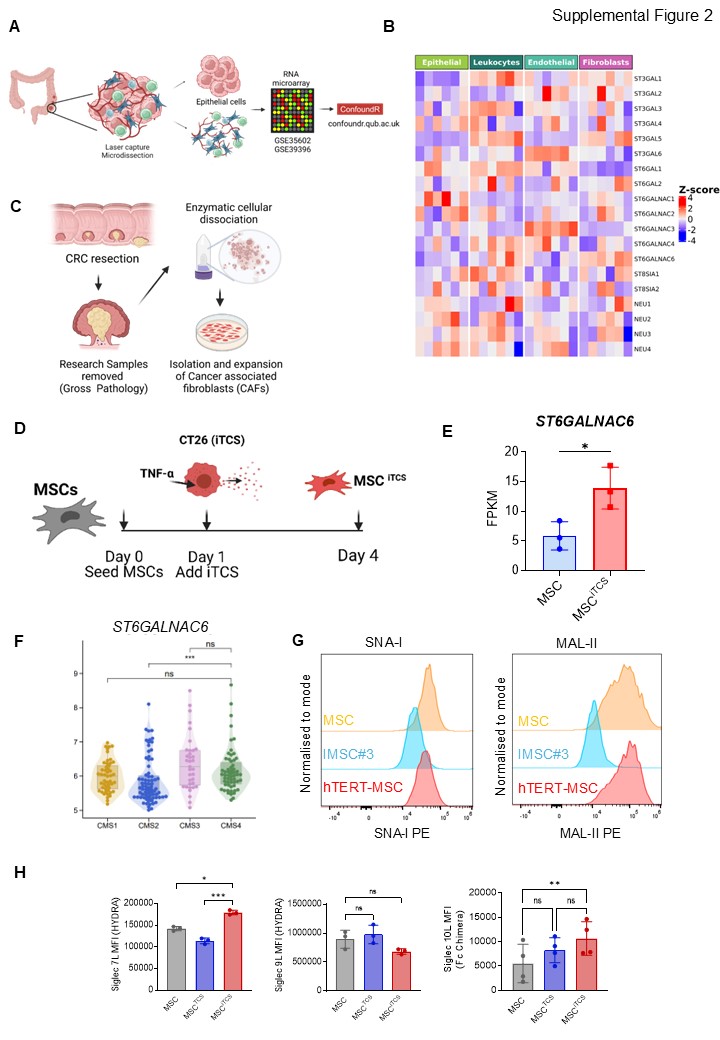

### Supplementary Figure 3

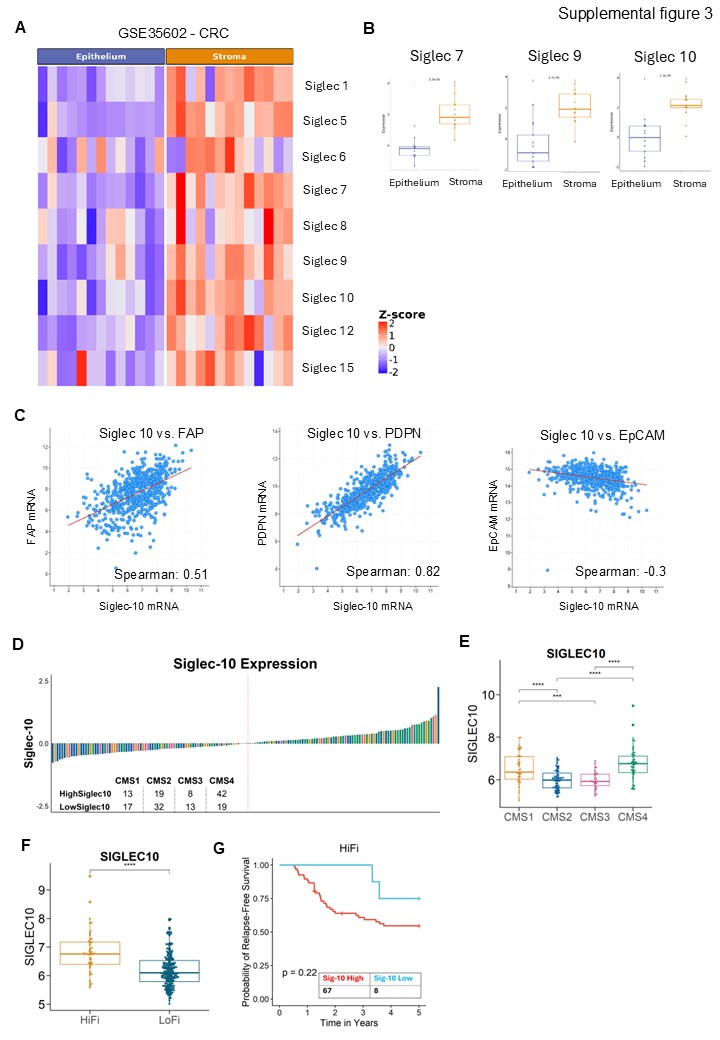

### Supplementary Figure 4

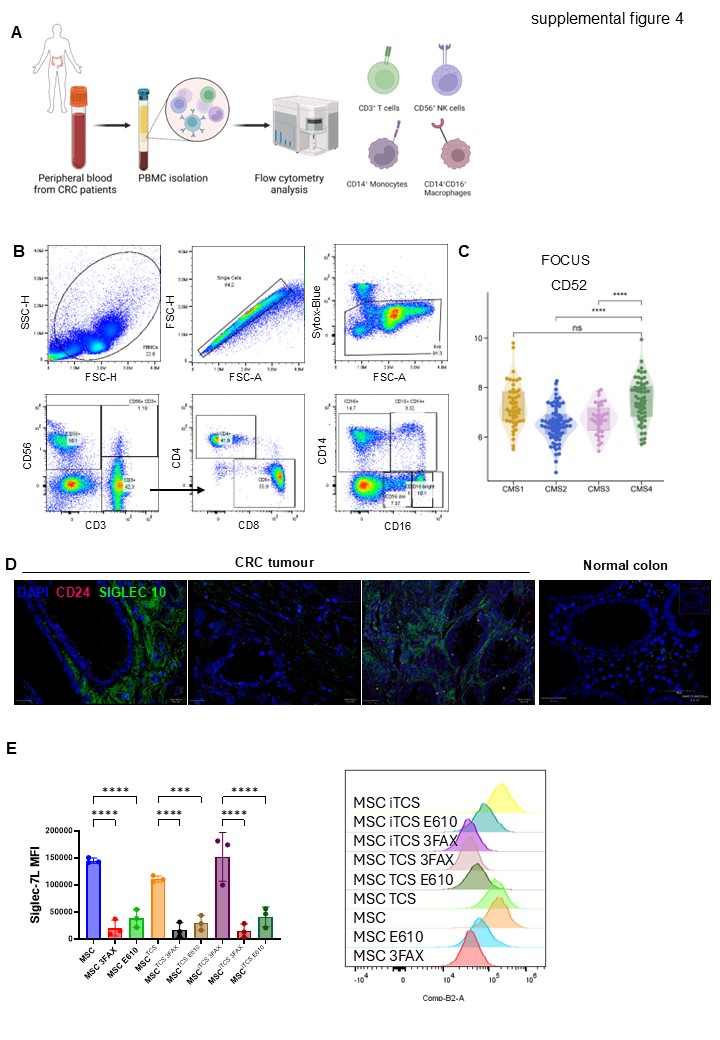

### Supplementary Figure 5

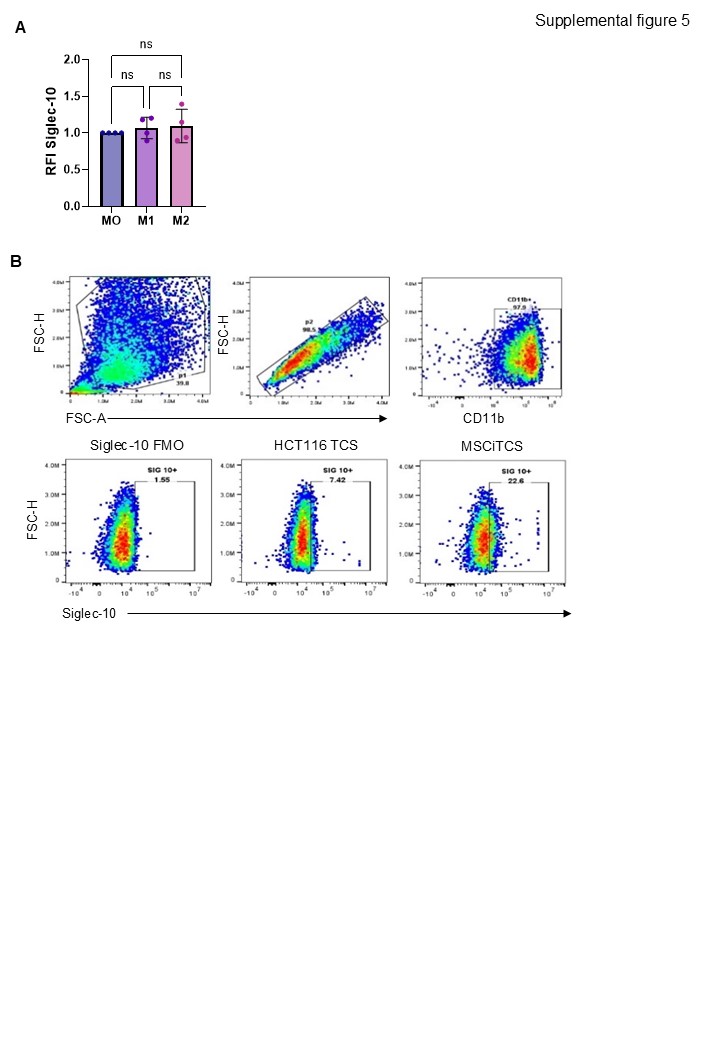

### Supplementray Figure 1

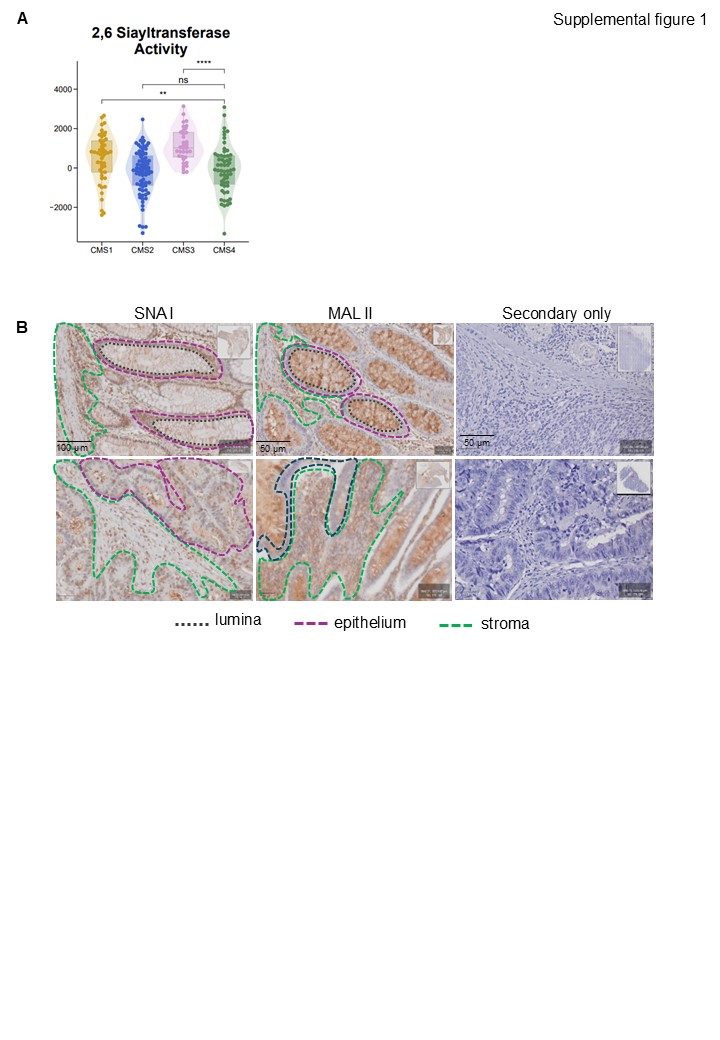
